# Breast cancer interactions with osteoclasts generate osteoclast-tumor hybrid-like cells through dynamic non-canonical cell fusion and cell-in-cell processes

**DOI:** 10.64898/2026.04.05.716538

**Authors:** King Hoo Lim, Damrongrat Siriwanna, Xining Li, Eunice Dotse, Meijun Wang, Choa Mun, Yusong Li, Xin Wang, Kwan Ting Chow

## Abstract

Macrophages/osteoclasts are highly fusogenic cells that interact closely with bone-metastatic breast cancer cells. These cancer cells adapt to bone microenvironments by undergoing osteomimicry, acquiring bone-like phenotypes. Exploration using human breast cancer-bone metastases dataset revealed that a small population of epithelial breast cancer cells express osteoclast-like and osteomimicry genes at the single-cell level. Cell fusion and cell-in-cell (CIC) processes are two uncommon yet prognostically significant mechanisms in cancer. We showed that co-culture between murine breast cancer cells and osteoclasts yielded a unique osteoclast phenotype through dynamic cell-in-cell (CIC) interactions and fusion-like behaviours between pre-osteoclasts/mature osteoclasts and breast tumor cells, resulting in osteoclast–tumor hybrid-like cells. These tumor cell interactions characterized by membrane retention and nuclear adjacency to host nuclei were consistently observed throughout osteoclast differentiation. Single-cell sequencing analysis and interpretative assays on hybrid-like cells revealed altered extracellular matrix (ECM) modification processes, immunoregulatory, and cancer-associated pathways compared to unfused osteoclasts. Tumor cells co-cultured with osteoclasts expressed hematopoietic and osteoclast-lineage factors more strongly than tumor cells cultured alone with their effects amplified under direct cell-cell contact. The presence of these hybrid-like cells was validated in human breast cancer-bone metastases. We propose that disseminated bone-tropic breast cancer cells were stimulated by osteoclasts to undergo a non-canonical, dynamic osteoclast differentiation and CIC formation to form hybrid-like cells that may facilitate bone metastatic lesions.

## Introduction

Cell fusion in cancer is indicated as a source for tumor heterogeneity[1, 2]. Particularly, heterotypic fusion between tumor cells and surrounding stromal cells can generate highly aggressive and invasive tumor variants [3, 4]. CIC formation is another significant cell interaction linked to cancer progression. It involves the engulfment of a cell by another through processes such as entosis, emperipolesis, cannibalism, and phagocytosis. Tumor cells can partake in both ends of the process where they exploit these CIC structures to evade immune cytotoxicity, gaining drug resistances, and acquiring aneuploidy. Despite their distinct mechanisms, the fundamentals on how cell fusion and CIC formation intertwine to influence cell fate decisions and contribute to tumor progression remain poorly understood.

Osteoclasts are multinucleated cells derived from cell fusion between myeloid precursors including macrophages and monocytes. While it is not known whether osteoclasts can form CIC structures, they have an intimate relationship with tumors exhibiting bone tropism and are recognized as the main culprit behind tumor-induced osteolysis, leading to bone destruction and skeletal-related complications[5]. Osteoclasts and tumors are known to influence each other through their respective secretome and extracellular vesicles, which create a vicious cycle promoting osteoclast bone resorption activity and tumor growth[6, 7]. Recently, direct cell-cell contact between osteoclasts and tumors has been suggested to hasten osteoclast maturation[8]. Furthermore, osteoclasts containing tumor cells within their cytoplasm have been reported[9]. These observations underline an underappreciated role of direct interactions between osteoclasts and tumors in bone metastatic development.

The occurrence of cell fusion/CIC formation between osteoclasts and tumors has been hinted in osteoclast-myeloma interactions in myeloma patients with bone metastases[10]. Osteoclasts in these patients were found to contain nuclei of B cell origins, where chromosomal translocations specific for myeloma cells were found within the osteoclast nuclei. Because of the propensity of most solid tumors to metastasize to the bone, we hypothesize that these tumors may undergo cell fusion/CIC formation with osteoclasts to form osteoclast-tumor hybrid-like cells.

In this study, we showed that a small subset of breast cancer cells from human bone metastases underwent transformation to express certain osteoclast-like/osteomimicry genes at a comparable level to canonical osteoclasts. We discovered that breast cancer and certain tumor types formed osteoclast-tumor hybrid-like cells via a dynamic combination of cell fusion and CIC formation processes. Tumor nuclei in these hybrid-like cells were transcriptionally active and gene expression profiling analyses in the hybrids uncovered enhanced tumor-promoting functions including angiogenesis, ECM-remodeling, motility, and the expression of cancer-associated pathways such as PI3K-AKT and Hippo signaling. Tumor cells in the presence of osteoclasts were induced to strongly express hematopoietic and osteoclast differentiation genes. We delved into how direct and paracrine interactions influenced osteoclasts and breast cancer cells and determined that close proximity and physical contact between the two cell types were crucial for their transformation. These close spatial association between osteoclasts and breast cancer cells were validated in-vivo. We identified cells that resembled our hybrid-like cells within the osteoclast population in human breast cancer-bone metastases. Taken together, we found that tumor cells were altered by osteoclasts to facilitate their fusion/CIC formation with developing osteoclasts. The resulting hybrid-like cells exhibited augmented osteoclast functions, particularly in ECM remodeling and angiogenesis while also expressing unique cancer-associated pathways. We propose that these osteoclast-tumor hybrid-like cells are a key player in bone metastatic lesions and may represent a novel target for preventing or intercepting bone metastasis.

## Results

### Rare breast cancer subpopulation in bone metastases exhibit osteoclast-like and osteomimicry gene signatures

Breast cancer cells with strong bone tropism have been described to adopt osteoclast and osteoblast-like gene expression via osteomimicry, with prevalence towards the latter[11, 12]. To investigate whether breast cancer cells can potentially mimic osteoclast functions, we sought to determine whether breast cancer cells in human bone metastases exhibit transcriptional signatures indicative of osteoclast-like and osteomimicry programs. We analysed a publicly available single-cell RNA sequencing dataset derived from ER+ breast cancer bone metastases (GSE190772). We depicted the various clusters containing epithelial and osteoclast populations defined by the original authors’ clustering (Fig. 1a). We curated an osteoclast (OC) / osteomimicry (OM) gene panel encompassing osteoclast-related markers (*Ctsk, Acp5, Ca2, Mmp2, Mmp9, Mmp14*) and osteomimicry-associated genes (*Spp1, Ibsp*) and compared their expression between epithelial and osteoclast clusters (Fig. 1b). As expected, global expression of these genes was predominantly confined to osteoclasts, excluding *Mmp2* and *Ibsp* as they are more prevalent in osteoblasts. However, we observed a small tail of epithelial cells exhibiting expression levels comparable to those of osteoclasts when we computed module scores for this OC/OM gene panel using AddModuleScore (Fig. 1c). This suggests the presence of a rare epithelial subset with osteoclast-like transcriptional features.

**Fig. 1:**
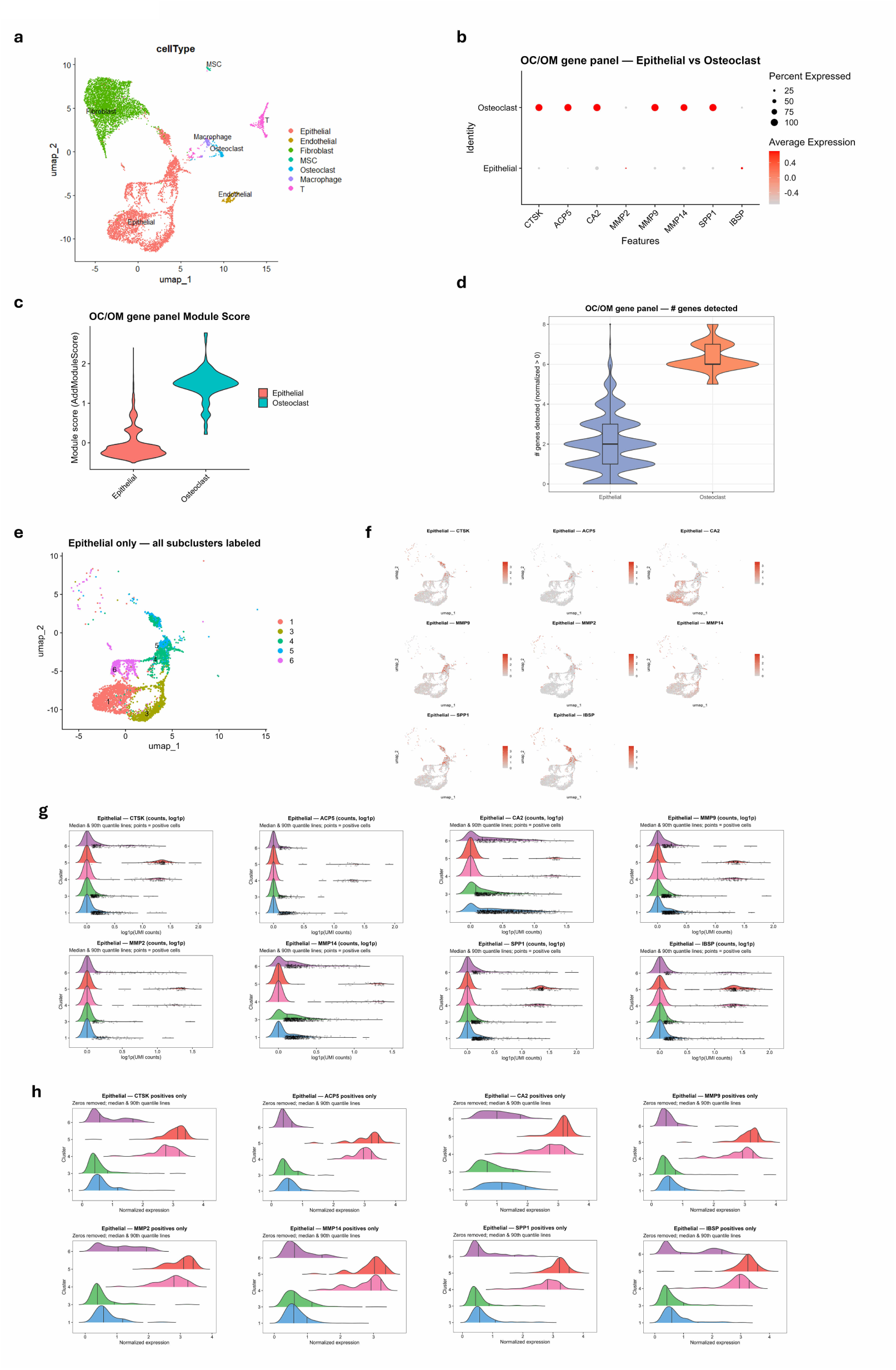
Osteoclasts fuse with various solid tumors to form osteoclast-tumor hybrids *in vitro*. **a**, Schematic diagram showing co-culture fusion assay between osteoclast precursors and solid tumor cell lines. **b**, Representative live cell images of an osteoclast-tumor hybrid (yellow outline) formed between RAW264.7-GFP osteoclasts (green) and 4T1-H2B-mRFP breast cancer cells (red). Tumor nuclei expressed H2B-mRFP (white arrow) within osteoclast cytoplasm. Scale bar, 100 μm. **c**, Schematic representation for Cre reporter lentiviral plasmid construct containing a floxed DsRed-E2 (stop cassette), followed by a downstream GFP. Cre mediated recombination excises the stop cassette and allows GFP to be expressed. **d,e**, Representative immunofluorescence images of BMM-derived osteoclasts co-cultured with EO771 Cre reporter. (**d**) Primary osteoclast-tumor hybrids (BMO Hybrid 1 & 2) (yellow outline) were formed in the co-culture assay with tumor nuclei identified based on RFP (DsRed-E2) fluorescence and round nuclei morphology (red arrows). (**e**) Co-cultured primary cells osteoclasts and tumor cells expressed Cre-recombinant product, GFP. Scale bars, 100 μm. **f**, Representative image showing a developing RAW264.7-GFP osteoclast-4T1-H2B-mRFP tumor hybrid surrounded by pre-osteoclasts and tumor cells. Tumor cells and pre-osteoclasts that were adjacent to the hybrid (blue arrows) had round morphologies while tumor cells located further away from the hybrid were epithelial-shaped (yellow arrow). **g-i**, Fluorescent time-lapse video showing cell fusion events between RAW264.7-GFP and 4T1-H2B-mRFP at various stages of osteoclast differentiation. (**g**) Tumor cell (yellow arrow) converged with other pre-osteoclasts to form a multinucleated osteoclast. Video stills were acquired from Movie S1 (**h**) Two osteoclast-tumor hybrids (yellow arrow) fused with each other. Video stills were acquired from Movie S2. (**i**) Individual tumor cells (yellow arrow) and pre-osteoclasts (white arrow) fused into an existing hybrid cell. Video stills were acquired from Movie S3.

To further quantify this phenomenon, we assessed the number of OC/OM genes detected per cell across both populations (Fig. 1d). While most epithelial cells expressed few or none of these genes, a minor fraction expressed five or more OC/OM genes at an equal level to osteoclasts. These findings indicate that breast cancer cells within bone metastases can acquire osteoclast-like and osteomimicry gene expression through some unknown mechanisms within the bone niche.

Having established that a minor epithelial fraction exhibits OC/OM features, we next delineated these OC/OM-high cells within the epithelial compartment that contains five subclusters (1, 3, 4, 5, 6) (Fig. 1e). UMAP visualization revealed that while subclusters 1, 3, and 6 displayed broad but low-level expression of panel genes, the most intense OC/OM signals were concentrated in a small fraction of cells within subclusters 4 and 5 (Fig. 1f). To quantify these patterns, we compared expression intensity versus prevalence for OC/OM genes across epithelial subclusters (Fig. S1a). We showed that subclusters 4 and 5 consistently rank in the top intensity quantiles but exhibit lower prevalence across their respective cell populations for most genes. In contrast, subclusters 1, 3, and 6 display broader prevalence at muted intensity, having a widespread yet weak expression state.

We further assessed the distributional expression for OC/OM genes and showed that log1p raw counts for subclusters 1, 3, and 6 delineated within 0.5, whereas subclusters 4 and 5 contain a distinct high-expression tail with values >1.0, indicating the presence of rare high-expressers (Fig. 1g). Tightening the parameters to positive-expression cells only yielded an even clearer separation with normalized values of 2 – 4 in subclusters 4 and 5 compared to 0 – 2 in subclusters 1, 3, and 6 (Fig. 1h). Together, these results relay that OC/OM transcriptional activity in the epithelial compartment is concentrated in a small subset of high expresser cells that are spatially enriched within subclusters 4 and 5, rather than broadly distributed at high levels.

### Breast cancer cells undergo dynamic cell fusion and CIC structure formation with developing osteoclasts

Metastasized cancer cells are frequently found in close proximity to osteoclasts, leading to physical interactions that activate osteoclast differentiation[13]. Together with our results showing co-expression of OC/OM genes in breast cancer cells, we hypothesize that these interactions may induce breast cancer cells to activate osteoclast-like differentiation programs to fuse with osteoclasts. To test whether osteoclastogenic conditions stimulate cancer cells to fuse with pre-osteoclasts *in vitro*, we co-cultured RAW264.7-GFP macrophages/osteoclast precursors with the breast cancer cell line, 4T1, which had been transduced to stably express RFP-tagged histone 2B (H2B-mRFP) (Fig. 2a). We observed that tumor nuclei expressing RFP signals were found within the mature multinucleated RAW264.7-GFP osteoclasts developed in the co-cultures (Fig. 2b). In contrast, we could not observe this occurrence under co-culture conditions without osteoclastogenic cytokines (data not shown). For primary osteoclasts, we similarly observed 4T1 tumor nuclei (H2b-mRFP) inclusion within the cytoplasmic bounds of differentiated mature osteoclasts derived from primary murine bone marrow cells (Fig. 2c). These observations denote that tumor cells can penetrate and reside within osteoclasts, while the precise mechanism behind this process remains unclear.

**Fig. 2:**
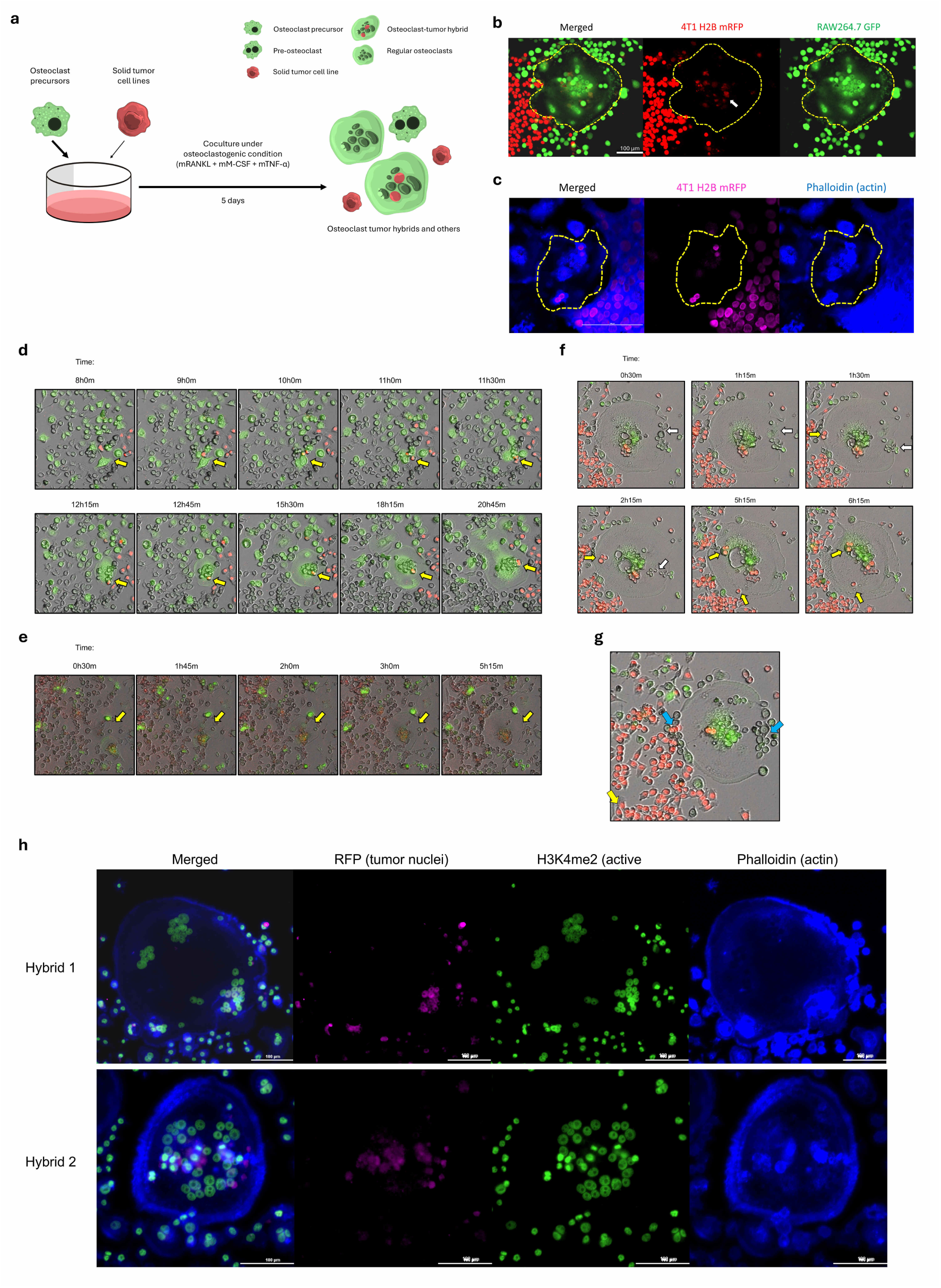
Osteoclasts fuse with various solid tumors to form osteoclast-tumor hybrids *in vitro*. **a**, Schematic diagram showing co-culture fusion assay between osteoclast precursors and solid tumor cell lines. **b**, Representative live cell images of a hybrid-like cell (yellow outline) formed between RAW264.7-GFP osteoclasts (green) and 4T1-H2B-mRFP breast cancer cells (red). Tumor nuclei expressed H2B-mRFP (white arrow) within osteoclast cytoplasm. Scale bar, 100 μm. **c**, Representative live cell images of a hybrid-like cell (yellow outline) formed between a primary bone-marrow-derived osteoclast (actin outline) and 4T1-H2B-mRFP breast cancer cells (magenta). Scale bar, 100 μm. **d-f**, Fluorescent time-lapse video showing cell fusion events between RAW264.7-GFP and 4T1-H2B-mRFP at various stages of osteoclast differentiation. (**d**) Tumor cell (yellow arrow) converged with other pre-osteoclasts to form a multinucleated osteoclast. Video stills were acquired from Movie S1 (**e**) Two osteoclast-tumor hybrids (yellow arrow) fused with each other. Video stills were acquired from Movie S2. (**f**) Individual tumor cells (yellow arrow) and pre-osteoclasts (white arrow) fused into an existing hybrid cell. Video stills were acquired from Movie S3. **g,** Live cell image showing RAW-246.7 GFP and 4T1-H2B-mRFP in distinct morphologies. Tumor and pre-osteoclast cells adjacent to an osteoclast displayed round cell shape (blue arrows) while tumor cells in growing colony were epithelial-shaped (yellow arrow). **h,** Representative immunofluorescent images for RAW264.7-GFP osteoclast-tumor hybrid-like cells stained for H3K4me2 transcriptional activity. Hybrids 1 and 2 contained tumor nuclei that fully colocalized with H3K4me2.

We then monitored the dynamics and cell movements by which tumor cells entered osteoclasts using a fluorescent live cell imager. During normal osteoclast development, individual mono-nucleated pre-osteoclasts fuse to form multinucleated mature osteoclasts. In co-cultures, we observed tumor cells integrating into developing osteoclasts that were in the process of fusing with other pre-osteoclasts (Fig. 2d, Movie S1). The tumor nuclei incorporated as one of the many nuclei within the osteoclasts that constitute a central nuclei system. Additionally, we observed fusion between multinucleated osteoclasts that already contained tumor nuclei (Fig. 2e, Movie S2). Upon merging of the cytoplasm, the osteoclast and tumor nuclei from each multinucleated osteoclast were observed to gather and converge together. Furthermore, we observed individual tumor cells and pre-osteoclasts entering a developed osteoclast with pre-existing tumor nuclei (Fig. 2f, Movie S3). Similarly, nuclei from these cells homed and gathered towards the central nuclei system. These observations were made in primary BM-derived osteoclasts co-cultured with EO771-DsRed-E2 as well (Fig. S2a,b, Movie S4).

To examine whether osteoclast-cancer fusion is limited to breast cancer, we extended our studies to B16F10 melanoma and CT26 colon cancer cell lines. We observed similar fusion hybrids formed between osteoclasts and these tumor lines (Fig. S3a,b). Collectively, these observations indicate that tumor cells gain access into osteoclasts through cell fusion that can occur at different stages throughout osteoclast differentiation. This phenomenon is also applicable to various solid tumor types other than breast cancer.

To delve into cellular dynamics, we observed that certain tumor and osteoclast nuclei within hybrid-like cells retained their cellular membranes as they entered or integrated into multinucleated osteoclasts (Fig. 2g). In contrast, some osteoclast nuclei lost their membrane integrity and merged completely into the shared cytoplasm. These findings suggest a combination of dynamic CIC structures and canonical osteoclast fusion mechanisms. Pre-osteoclasts are known to adopt a rounded morphology prior to osteoclast fusion[14]. Notably, both pre-osteoclast and tumor nuclei in our observation assume rounded shape before fusion or CIC entry. This aligns with reports that morphological transition precedes incorporation into multinucleated osteoclasts.

Osteoclast nuclei can be present as either in a viable state with intact nuclei or apoptotic condition having fragmentated patterns[15]. Additionally, osteoclasts can phagocytose apoptotic cells for debris clearance[16]. To discern the tumor nuclei state in our hybrid-like cells, we assessed the RFP+ tumor nuclei for H3K4me2 post-translational modification that marks actively transcribed loci[17]. Many of our observations resembled Hybrid 1, wherein the RFP and H3K4me2 signals colocalized in intact tumor nuclei within the osteoclasts, indicating that they were viable and transcriptionally active (Fig. 2h). However, we also noted the presence of fragmented, apoptotic tumor nuclei mixed in with viable ones in some hybrid-like cells such as the case for Hybrid 2. This suggests that while a subset of tumor nuclei integrated within osteoclasts remains transcriptionally active, others appear fragmented and likely non-functional. Importantly, these transcriptionally active tumor nuclei may contribute to gene expression in osteoclast-tumor fusion hybrid-like cells.

Focusing on the nuclear fusion process between tumor and osteoclast nuclei, we tracked the fate of tumor nuclei once they were incorporated into an osteoclast. While most tumor nuclei were retained adjacent to osteoclast nuclei as previously observed, we managed to capture instances where a tumor nucleus underwent shrinkage and subsequently disintegrated adjacent to osteoclast nuclei (Fig. S4a, Movie S5). These observations bear great resemblance to entosis as part of the cell-in-cell phenomenon where internalized cells are led to cell death by lysosomal degradation. Collectively, our findings indicate that breast cancer cells enter osteoclasts through a dynamic interplay of canonical osteoclast fusion and cell-in-cell structure formation. The functional impact of these hybrid-like cells is likely heterogeneous due to differences in integrity, transcriptional activity, and number of integrated tumor nuclei. Consequently, this variability may generate distinct phenotypes of osteoclast–tumor hybrid-like cells rather than a single uniform outcome.

### Osteoclast-tumor hybrid-like cells exhibit distinct functional profiles compared to non-hybrid osteoclasts

To delve into the identity and characteristics of the osteoclast-tumor hybrid-like cells, we devised a laser microdissection approach to precisely isolate these hybrids within the co-cultures (<u>o</u>steoclast <u>h</u>ybrid-like <u>c</u>ells; OHC) (Fig. 3a). As controls, we used the same approach to isolate osteoclasts without tumor nuclei in co-cultures (<u>o</u>steoclasts in <u>c</u>o-culture; OC), as well as non-hybrid osteoclasts in osteoclast-only cultures (<u>o</u>steoclasts; O). Corresponding cell types were imaged prior to isolation (Fig. 3b). Isolated cells with good RNA quality were subjected to downstream Smart-Seq 2 single-cell sequencing (Fig. S5a,b). Comparison of differentially expressed genes (DEGs) revealed that OHC and OC had substantial number of DEGs when compared against O, with OHC having twice as many DEGs than OC when compared against O (Fig. 3c). When comparing OHC and OC, there were close to 200 DEGs, indicating that these two cell types within the same co-culture were significantly different. We compared the normalized z-scores for DEGs across the three cell types in a heatmap, which showed clear differences in upregulated and downregulated genes in OHC as compared to OC and O (Fig. 3d). Meanwhile, the two non-hybrid osteoclast groups (OC and O) shared resemblance in their gene expression profiles. These results demonstrated that OHCs are a distinct and unique cell type from non-hybrid osteoclasts.

**Fig. 3:**
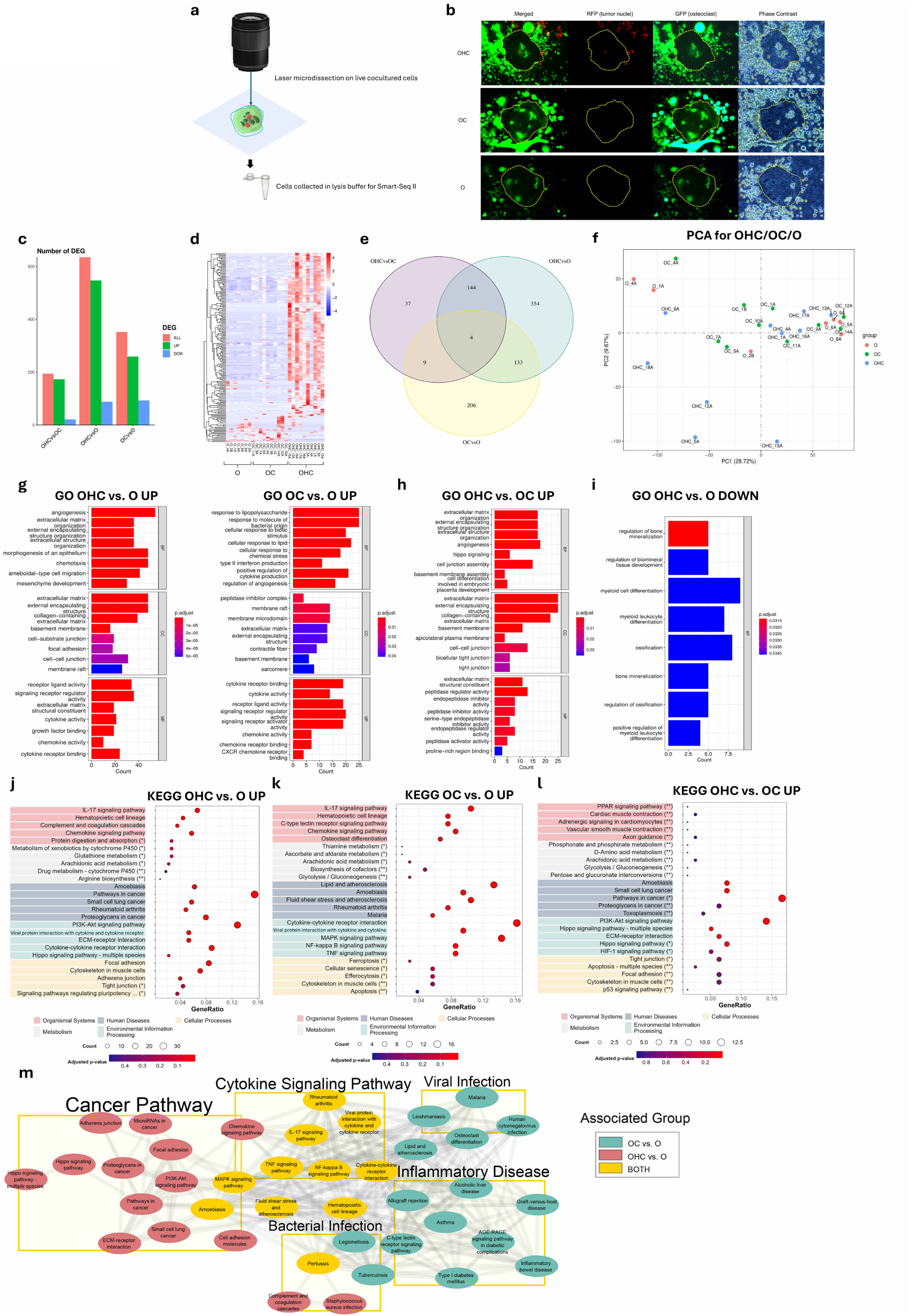
Osteoclast-tumor hybrids are transcriptionally distinct from non-hybrid osteoclasts. **a**, Schematic diagram displaying live cell laser microdissection protocol on RAW264.7-GFP osteoclasts co-cultured with 4T1-H2B-mRFP breast cancer cells to excise hybrids and other cells of interest for downstream single cell sequencing. **b**, Representative fluorescent images for three osteoclast sample types with hybrids (OHC) (n = 10), non-hybrid osteoclasts from co-culture (OC) (n = 10), and non-hybrid osteoclasts from osteoclast-only culture (O) (n = 7) with osteoclasts highlighted in yellow outlines. **c**, Bar graph showing three DEGs comparison groups with OHC vs. O, OC vs. O, and OHC vs. OC. Upregulated, downregulated, and overall DEGs were displayed for each group. **d**, Heatmap displaying both upregulated and downregulated DEGs amongst the three osteoclast sample types. **e**, Venn diagram showing overlapping and unique overall DEGs amongst the three osteoclast comparison groups. **f**, PCA analysis plot for individual samples from three osteoclast sample types. **g,h** Gene ontology (GO) analyses for hybrids and non-hybrid osteoclasts on three comparison groups (OHC vs. O, OC vs. O, and OHC vs. OC). Upregulated gene functions on biological processes (BP), cellular components (CC), and molecular functions (MF) for (**g**) OHC vs. O, OC vs. O, and (**h**) OHC vs. OC. (**i**) Attenuated gene functions on BP, CC, and MF for OHC vs. O. Gene functions and their respective gene counts were shown. p.adjusted value < 0.05. **j-l,** KEGG pathway analyses for the same comparison groups. Top five pathways augmented in various categories for (**j**) OHC vs. O, (**k**) OC vs. O, and (**l**) OHC vs. OC. Pathways were shown in order of significance with following annotations: no asterisk – p.adjusted value & p value < 0.05, one asterisk (*) – p.adjusted value > 0.05 & p value < 0.05, two asterisks (**) – p. adjusted value & p value > 0.05. (**m**) Network analysis for significant pathways enriched in OHC vs. O (red), OC vs. O (blue), and both (yellow). p.adjusted value < 0.05.

Honing in on the identity of the DEGs, OHC vs. O and OC vs. O had a substantial number of non-overlapping DEGs of 498 and 215 DEGs respectively (Fig. 3e). In contrast, the majority of DEGs in OHC vs. OC was also present in OHC vs. O, demonstrating that OHC are dissimilar to osteoclasts overall in terms of gene expression. Principle component analysis (PCA) showed that OC and O bore a high degree of resemblance with one another (Fig. 3f). Conversely, OHC showed a dispersed pattern with some forming a unique cluster while the others spread amongst the non-hybrid osteoclasts. Overall, these results indicate that osteoclast-tumor hybrid-like and non-hybrid osteoclasts within the same environment differed in gene expression from one another and from osteoclasts in the absence of tumor cells. However, we do note that OHC population had a more heterogeneous expression, which corroborates with previously discussed tumor nuclei transcriptional activity in hybrid-like cells.

We delved into the gene functions enriched in OHC by analyzing the gene ontology (GO) of DEGs in OHC vs. O and OC vs. O. (Fig. 3g). We found enriched biological processes (BP) including angiogenesis, ECM-remodeling, cell motility, and epithelial/mesenchymal transformation in OHC vs. O. Moreover, notable cellular components (CC) like focal adhesion and cell-cell junctions were enhanced. On the other hand, OC vs. O exhibited abundant genes associated with cellular and inflammatory responses like type II interferons and other cytokine production. Enriched components in the CC category in OC vs. O displayed mostly membrane-related processes. Both OHC vs. O and OC vs. O groups displayed enhanced cytokine and chemokine signalling under molecular functions (MF).

As expected, the enriched functions in OHC versus OC largely overlapped with those observed in OHC versus O (Fig. 3h). Notably, OHCs exhibited strong upregulation of programs related to extracellular matrix organization, basement membrane assembly, and cell–cell/tight junction formation, alongside Hippo signaling and angiogenesis. These features indicate a shift toward matrix remodeling and niche construction rather than classical resorptive osteoclast activity.

Elevated peptidase regulator and inhibitor activity further suggests controlled proteolysis, favoring structured ECM deposition. Downregulated OHC vs. O gene functions aligned with the phenotype shift, wherein it showed decline for functions governing bone mineralization, ossification, and myeloid differentiation (Fig. 3i). These point towards the loss of classical osteoclast maturation and mineral-degrading capacity. Collectively, these findings support that hybrid-like cells adopt a stromagenic, adhesion-oriented, and pro-angiogenic phenotype, consistent with epithelial-like traits and tumor-supportive functions, whereas non-hybrid osteoclasts remain more skewed toward pro-inflammatory responses in the presence of tumor cells.

Kyoto Encyclopedia of Genes and Genomes (KEGG) pathway analyses for corresponding comparisons corroborated most of the differences observed in GO (Fig. 3j,k,l). OHC vs. O displayed activation of PI3K–Akt and Hippo signaling, ECM–receptor interaction, focal adhesion, and junctional pathways, alongside cytokine/chemokine cascades and complement signaling (Fig. 3j). These changes indicate reprogramming toward an inflammatory, mechanosensitive, adhesion-rich, pro-survival state that remodels matrix, supports angiogenesis, and may confer drug tolerance through enhanced xenobiotic and glutathione metabolism. In contrast, OC vs. O upregulated immune and inflammatory signaling (IL-17, chemokine, C-type lectin receptor), hematopoietic lineage, and osteoclast differentiation pathways, alongside NF-κB/MAPK/TNF signaling and cytokine–cytokine interactions (Fig. 3k). These enrichments reflect a pro-inflammatory, osteoclast-activating state with stress-response and apoptotic programs, contrasting with the stromagenic phenotype observed in hybrid-like cells.

In OHC vs. OC, we observed that OHC had higher enrichment on aforementioned cancer-associated signalling including PI3K-AKT and Hippo signalling pathways and ECM-remodelling functions (Fig. 3l). A KEGG pathway network analysis further substantiated the trends observed in GO (Fig. 3m). OHC vs. O was enriched with pathways contributing to cancer progression while OC vs. O emphasized on immune response and inflammatory disease pathways. The two comparisons shared enrichment in several cytokine signalling pathways. Together, these outcomes underscore that hybrid-like cells prioritize cancer-promoting and adhesion-driven pathways, whereas co-cultured osteoclasts predominantly engage on immune and inflammatory programs.

### Breast cancer cells express unique genes in the presence of osteoclasts

The significant changes observed in osteoclasts within co-cultures prompted us to investigate how the presence of osteoclasts altered the phenotype of tumor cells. We isolated tumor cells from co-cultures (T_CO) using laser microdissection for single-cell sequencing to explore their gene expression profile (Fig. 4a). Controls included tumor cells treated with osteoclastogenic cytokines (T_T) and untreated tumor cells (T_UT). For T_CO, we took precautions to isolate tumor cells within its own clusters to avoid contamination with osteoclasts. Prior to sequencing, we performed quality control on isolated RNA (Fig. S6a,b).

**Fig. 4:**
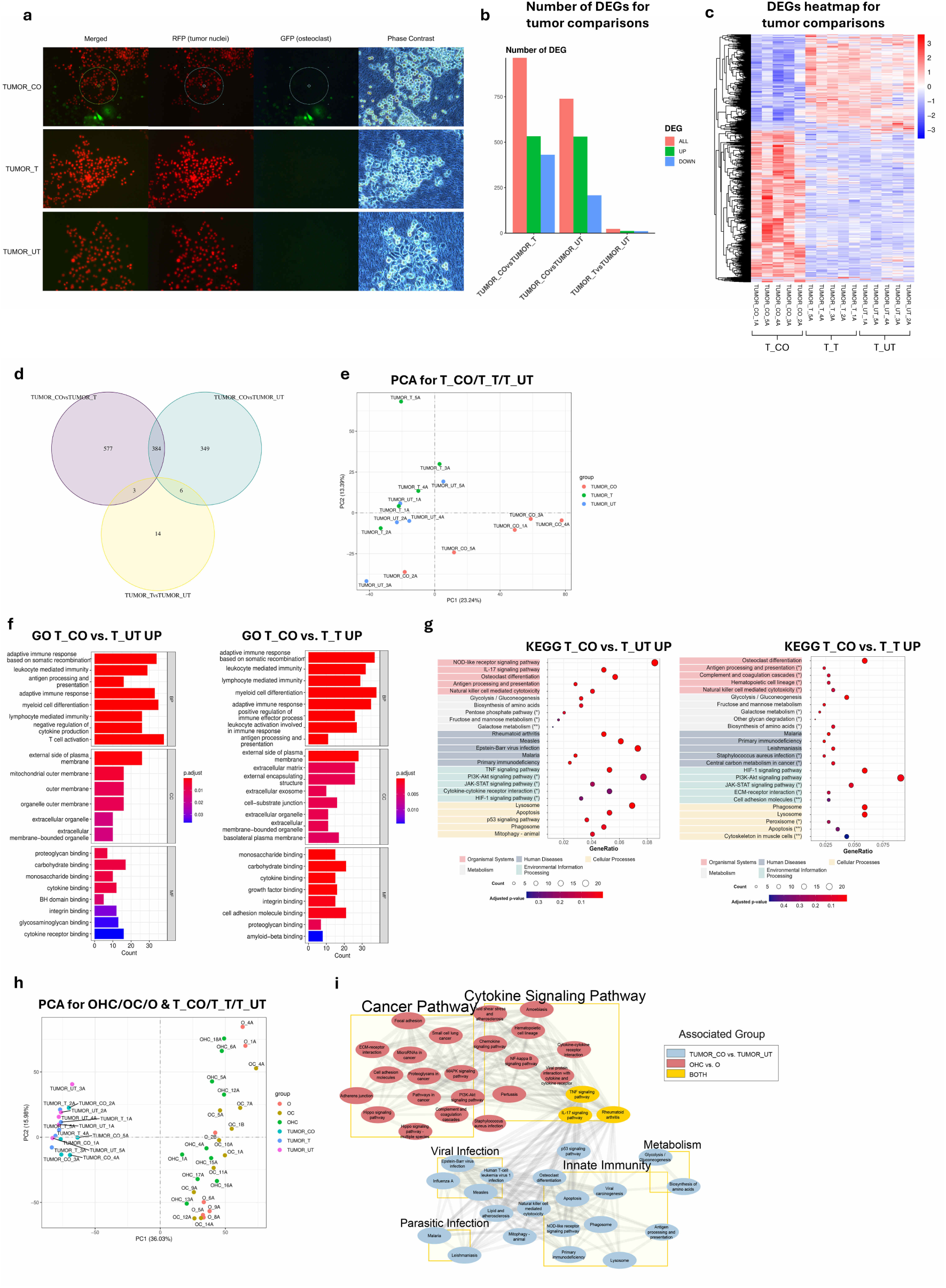
Breast cancer cells express unique gene expression profile in the presence of osteoclasts. **a**, Representative fluorescent images for three tumor sample types with tumors from co-culture (TUMOR_CO) (T_CO), osteoclastogenic cytokine-treated tumors (TUMOR_T) (T_T), untreated tumors (TUMOR_UT) (T_UT). Example for T_CO isolated cells were highlighted in circle. **b**, Bar graph showing three DEGs comparison groups with TUMOR_CO vs. TUMOR_T, TUMOR_CO vs. TUMOR_UT, and TUMOR_T vs. TUMOR_UT. **c**, Heatmap containing both upregulated and downregulated DEGs amongst the three tumor sample types. **d**, Venn diagram depicting overlapping and unique overall DEGs amongst the three tumor comparison groups. **e**, PCA analysis plots for individual tumors from three tumor sample types. **f**, GO analyses for tumor samples with three comparison groups (T_CO vs. T_UT and T_CO vs. T_T). Upregulated gene functions on BP, CC, and MF for T_CO vs. T_UT, and T_CO vs. T_T. **g**, KEGG pathway analyses for the same tumor comparison groups. Top five pathways augmented in various categories for T_CO vs. T_UT and T_CO vs. T_T. Pathways were shown in order of significance with following annotations: no asterisk – p.adjusted value & p value < 0.05, one asterisk (*) – p.adjusted value > 0.05 & p value < 0.05, two asterisks (**) – p. adjusted value & p value > 0.05. **h**, PCA analysis plots for individual samples from both osteoclast and tumor sample types. **i**, Network analysis showing significant pathways enriched in T_CO vs. T_UT (light blue), OHC vs. O (red), and both (yellow). p.adjusted value < 0.05.

Comparing T_CO with T_T and T_UT controls, we found that T_CO had a large number of DEGs, indicating a significant change in gene expression in tumor cells co-cultured with osteoclasts (Fig. 4b). In contrast, treatment with osteoclastogenic cytokines (T_T) did not result in notable changes as compared to untreated tumor cells (T_UT). T_CO expressed distinct DEGs while the two controls, T_T and T_UT resembled each other (Fig. 4c).

Examining the DEGs revealed that T_CO vs. T_UT and T_CO vs. T_T shared approximately half of their DEGs (Fig. 4d). Conversely, T_T vs. T_UT had only a handful of overlapping DEGs with T_CO vs. T_UT and T_CO vs. T_T. PCA analysis showed that T_CO formed its own cluster that was clearly distinct from T_T and T_UT, which largely grouped together (Fig. 4e). These highlight a unique gene expression program in T_CO compared to the other two tumor groups. Overall, tumors in co-culture underwent significant alterations in their gene expression as compared to cytokine-treated and untreated tumor controls, which resembled to each other. This ultimately pinpoints that osteoclastogenic cytokines alone are insufficient to induce considerable changes in tumors and that the physical presence of osteoclasts is necessary for significant gene expression alterations in tumors.

To dissect how T_CO were functionally altered, we performed GO analyses for upregulated DEGs in T_CO vs. T_UT and T_CO vs. T_T (Fig. 4f). We found T_CO exhibited elevated expression in genes correlated with hematopoietic cell differentiation and antigen processing functions. Simultaneously, extracellular and membrane functions along with ligand-receptor activities were enhanced. These changes suggested that in the presence of osteoclasts, tumor cells were induced to undergo cellular differentiation, which may involve outer membrane stimuli and ligand-receptor activities. We delved further into individual DEGs and identified notable upregulated genes such as *Dcstamp, Ocstamp*, *Oscar*, and *Trem2*, all of which are essential for osteoclast fusion and differentiation (Fig. S7a,b).

Similar trends were corroborated in KEGG analyses for T_CO vs. T_UT and T_CO vs. T_T (Fig. 4g). In both comparisons, immune response pathways and osteoclast differentiation were amongst the highly enriched pathways. Along the same vein, several enhanced cellular processes included phagosome and lysosome regulation, which are commonly associated with immune functions. Interestingly, we observed PI3K-AKT signalling to be significantly enhanced in T_CO vs. T_T, which was also found to be enhanced in OHC. But unlike the hybrids, T_CO have significantly enhanced glycolysis amongst other metabolism pathways. Further, we found that protein translation functions in T_CO vs. T_T were highly attenuated in both GO and KEGG analyses (Fig. S8a,b). Overall, gene function and pathway alterations in T_CO indicate that breast cancer cells were stimulated by the presence of osteoclasts to undergo osteoclast-like differentiation accompanied with activation of various hematopoietic differentiation and antigen presentation processes.

Noting potentially similar enriched pathways that were shared between tumor in co-culture (T_CO) and hybrid-like cells (OHC), we compiled a universal PCA analysis including both tumor and osteoclast cell groups (Fig. 4h). There was a clear separation between osteoclasts and tumor cells regardless of treatment, indicating that the two cell groups had inherently distinct gene expression profiles. Importantly, the hybrids resembled osteoclasts to a higher degree than to tumors. KEGG pathway network analysis uncovered shared and distinct pathways between OHC vs. O and T_CO vs. T_UT (Fig. 4i). Hybrid-like cells showed increased cancer-related and cytokine signaling, while co-cultured tumors exhibited metabolic and hematopoietic pathway activation. IL-17, TNF, and RA signaling were among the few pathways commonly upregulated in both hybrid-like cells and tumors. Our results demonstrate that tumor cells in co-culture assume a unique cellular identity as they exhibit predominantly distinct functions when compared to hybrid-like cells.

### Spatial association with osteoclasts influences breast cancer cell behaviour and microenvironment

Previous studies have demonstrated that tumor cells can physically interact with osteoclasts and osteoblasts in vivo, highlighting the importance of spatial association[13, 18, 19]. In our model, breast cancer cells adopted rounded morphologies before integrating into osteoclasts through fusion and cell-in-cell processes, suggesting that proximity may influence tumor adaptation. To investigate this, we compared direct co-culture with indirect transwell co-culture (Fig. 5a). We collected supernatants from both conditions for cytokine array assessment (Fig. 5b). Cytokine profiling on supernatants from the two co-culture designs revealed distinct patterns with direct association promoted immunosuppressive and pro-angiogenic factors such as CCL22, IL-10, VEGFA, and PDGFb, whereas paracrine-only interaction induced higher levels of pro-inflammatory cytokines including CSF2, IL-23, CXCL16, and others. Direct contact also increased mediators linked to extracellular matrix remodeling, including MMP2, MMP9, and PTX3, which contribute to niche construction and angiogenesis. These findings indicate that close spatial association between osteoclasts and breast cancer cells shift the immediate microenvironment towards an immunosuppressive, angiogenic, and matrix-remodeling phenotype, contrasting with the pro-inflammatory profile observed under paracrine signaling alone.

**Fig. 5:**
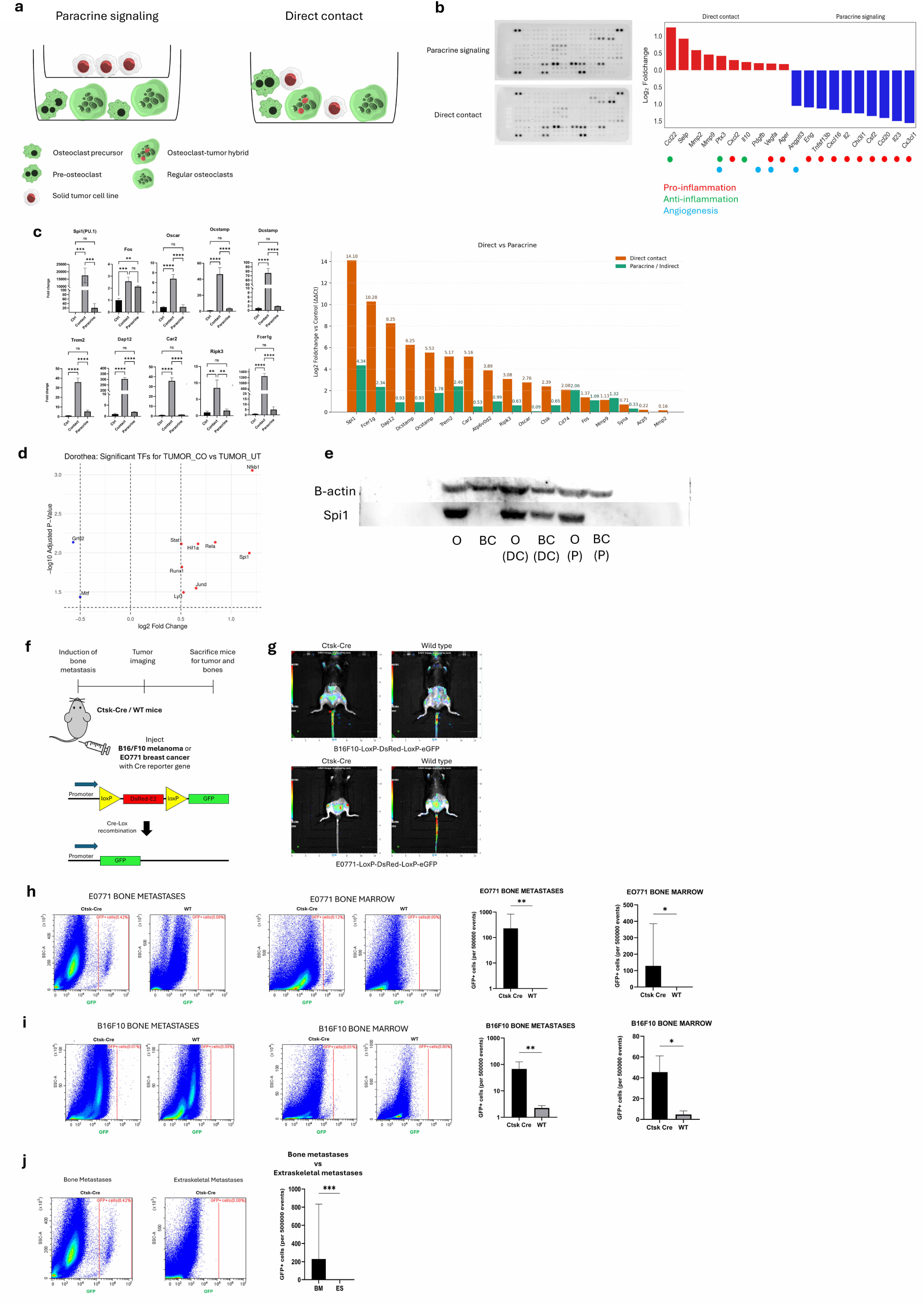
Direct contact with osteoclasts reprograms breast cancer cell transcriptome. **a,** Schematic diagram showing two co-culture setups for RAW264.7 preosteoclasts and 4T1 breast cancer cells under indirect (paracrine) and direct contacts. **b,** Cytokine array membrane blots for the two conditions. Top ten cytokine factors (fold change) enhanced in either direct contact or paracrine interaction conditions compared to each other. Cytokines associated with pro-inflammation (red), anti-inflammation (green), and angiogenesis (blue) were labelled. No significance was shown for cytokine array data. **c,** qPCR analysis on 4T1 isolated from direct contact and paracrine conditions with 4T1 from regular culture as baseline control. **d,** Dorothea analysis on TUMOR_CO vs TUMOR_UT showing differentially activated transcription factors. **e,** Western blot analysis for RAW264.7 osteoclasts and 4T1 cells under various culture conditions: osteoclasts only - O, breast cancer cells only - BC, osteoclasts from direct co-culture - O (DC), breast cancer cells from direct co-culture - BC (DC), osteoclasts from paracrine co-culture – O (P), breast cancer cells from paracrine co-culture – BC (P). **f,** Schematic diagram depicting bone metastasis induction on Ctsk-Cre and wild type (WT) mice models using EO771 breast cancer and B16-F10 melanoma expressing DsRed-E2-to-GFP Cre reporter system. **g,** Fluorescent images for DsRed-E2 fluorescent detection on mice with successful bone metastasis induction using EO771 and B16F10 expressing Cre reporter. **h,i,** Flow cytometry analysis on gated single cells showing DsRed-E2+ and GFP+ cells in EO771 and B16-F10 bone metastases derived from Ctsk-Cre mice. Statistics for GFP+ Cre recombined cells (per 500000 events) for EO771 bone metastases in Ctsk-Cre (n = 8) & WT (n = 5), and EO771 bone marrow in Ctsk-Cre (n = 5) & WT (n = 4). Statistics for GFP+ Cre recombined cells (per 500000 events) for B16F10 bone metastases in Ctsk-Cre (n = 6) & WT (n = 4), and B16F10 bone marrow in Ctsk-Cre (n = 5) & WT (n = 3). **j**, GFP+ recombined cells in bone metastases versus extraskeletal metastases for EO771 model. Statistics for GFP+ Cre recombined cells (per 500000 events) in EO771 with bone metastases (n = 8) & extraskeletal metastases (n = 6). Data are shown as means ± s.d. *P < 0.05; **P < 0.01; ***P < 0.001.

To determine whether proximity and interaction with osteoclasts drive osteoclast-like reprogramming in tumor cells, we analyzed gene expression in breast cancer cells recovered from direct and indirect co-cultures. Our qPCR data revealed a striking induction of osteoclast and hematopoietic programs under direct contact (Fig. 5c). Spi1 showed the most pronounced change, with approximately 14 log₂ fold increase compared to no co-culture control, whereas paracrine interaction induced only minimal changes. Other genes involved in osteoclast signaling and fusion, including Fcer1g, Dap12, Dcstamp, Ocstamp, and Trem2, were similarly elevated under direct contact. Functional osteoclast regulators such as Car2 and Atp6v0d2 also showed strong induction, consistent with enhanced acidification and fusion capacity. Notably, several genes from our OC/OM panel such as Car2, Ctsk, Acp5, Mmp9, and Mmp2 were upregulated in tumors under direct association, consistent with our earlier single-cell data mining results that these markers define a rare subset of breast cancer cells in bone metastases.

To substantiate these observations, we performed Dorothea analysis on our own Smart-seq2 single-cell data comparing TUMOR_CO (direct co-culture) and TUMOR_UT (no co-culture control). Spi1 emerged as one of the significantly upregulated transcription factors, alongside Nfkb1, Rela, and Hif1a (Fig. 5d). This reinforces the activation of hematopoietic and osteoclast-related transcriptional programs in tumor cells in the presence of osteoclasts.

We next examined protein-level changes by Western blot detection on Spi1 and found that it was detected in all CD45⁺ osteoclasts across conditions, as expected. However, it was expressed only in breast cancer cells under direct contact and not in paracrine or no co-culture control conditions (Fig. 5e). This confirms that spatial association with osteoclasts induces hematopoietic-like traits in tumor cells at both RNA and protein levels.

To validate that osteoclast-tumor interactions occur *in vivo*, we utilized the Ctsk-Cre mouse model, where the expression of Cre recombinase is largely restricted to osteoclasts. To establish a bone metastasis model, we injected breast cancer (EO771) or melanoma (B16F10) cells expressing the Cre reporter gene (lox-DsRed-STOP-lox-GFP) into the caudal artery of the mice (Fig. 5f). Both breast cancer and melanoma frequently metastasize to bones and hence were used as bone metastatic tumors for our investigation. We monitored the development of bone metastases via DsRed fluorescence (tumor cells) using *in vivo* animal imaging (Fig. 5g). At the end point, we collected bone metastatic tissues to assess the presence of tumor cells with intact Cre reporter (DsRed+) and Cre-recombined tumor cells (GFP+). These GFP+ tumor cells were denoted as osteoclast-associated tumors. Flow cytometry of bone metastatic lesions and bone marrow revealed GFP+ cells exclusively in Ctsk-Cre mice with bone metastases, but not in wild-type controls for both EO771 and B16F10 models (Fig. 5h,i). Gating strategies for acquiring single cells for analysis were shown (Fig. S9a). In addition, we also assessed for GFP+ cells in tumors that have developed in extraskeletal sites, including the liver, spleen, lungs, and the abdominal area. Analysis of extraskeletal metastases showed no GFP+ cells, indicating that Cre recombination occurred specifically within the bone metastatic niche (Fig. 5j). These findings demonstrate that tumors physically associate with Ctsk⁺ osteoclast lineage cells in bone metastases, supporting the existence of osteoclast-associated tumors. In a nutshell, our results show that tumors in bone metastatic lesions form close physical associations with osteoclasts, and these interactions likely reprogram tumor cells to adopt osteoclast-like traits and activate matrix-remodeling programs.

### Osteoclast-tumor hybrid-like cells and tumors from co-culture are found in human breast cancer-bone metastasis

To validate the presence of these hybrid-like cells in human patients, we inspected a single cell sequencing dataset that analyzed bone metastases collected from a patient diagnosed with estrogen receptor+ (ER+) breast cancer[20]. In the dataset, we utilized pre-defined cell clusters and labels provided by the authors (Fig. 6a). We focused on the osteoclast population and utilized gene set variation analysis (GSVA) to identify cells that resembled our OHC and OC based on significant DEGs derived from OHC vs. O and OC vs. O comparisons respectively (Fig. 6b). We ranked the cells based on their GSVA similarity score and labelled the top ranked cells for OHC and OC correspondingly. Interestingly, we found that cells with the highest similarity to OHC formed an isolated cluster of its own. In contrast, OC-like cells were distributed in two regions and partially coincided with OHC-like cells. This trend corroborates with our Smart-Seq 2 analysis where OHC and OC shared partial similarities in the PCA and network analysis.

**Fig. 6:**
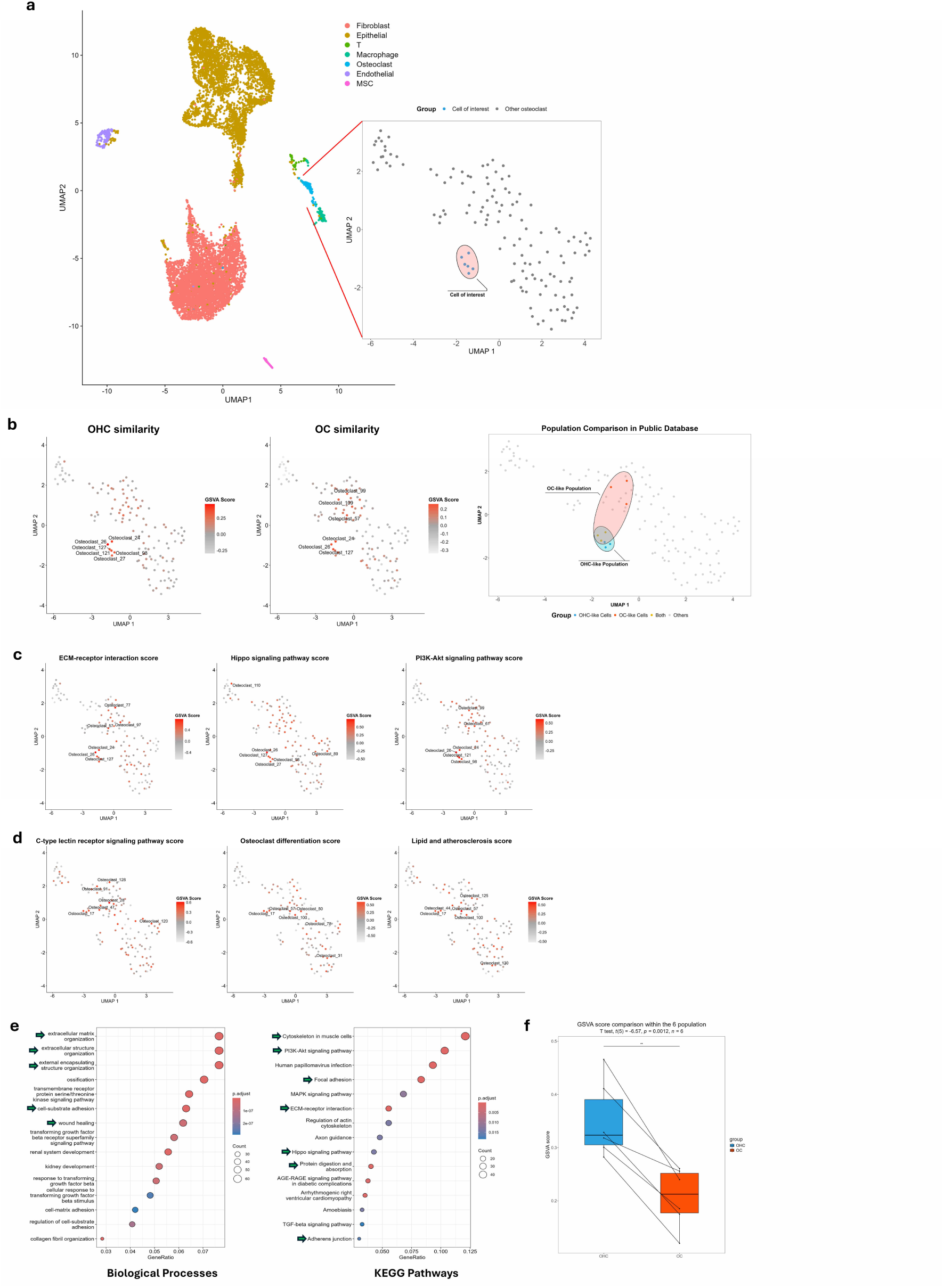
Human ER+ breast cancer-bone metastasis contains cells resembling osteoclast-tumor hybrids and tumors from co-culture. **a,** Cells of interest resembling hybrid-like cells in osteoclast population pre-defined by authors from GSE190772 containing scRNA data derived from ER+ human patient breast cancer-bone metastasis. **b,** Gene set variation analyses (GSVA) for overall DEGs derived from OHC vs. O and OC vs. O mapped and compared against osteoclast population. Similarity scores for OHC vs. O and OC vs. O plotted across osteoclast population with top six scores labelled on individual osteoclasts. Individual osteoclast cells in GSVA with scores > 0 were highlighted in ascending red intensity. Expected OHC- (red) and OC-like (blue) populations were interpolated as highlighted clusters within osteoclast population. **c,d,** GSVA for several unique upregulated pathways in (**c**) OHC vs. O and (**d**) OC vs. O respectively. **e,** GO (BP) and KEGG pathway breakdown on OHC-like cluster with several highlighted functions and pathways resembling hybrid-like cells in our data (green arrows). **f,** GSVA score comparison for OHC vs. O and OC vs. O centroids for OHC-like cell cluster. Statistical analysis via paired T test (n = 6); p = 0.0012.

To gain insights into the phenotypes of the two cell types, we evaluated GSVA similarity for several uniquely upregulated pathways in OHC vs. O and OC vs. O comparisons (Figure 6c,d). We found that cancer-associated pathways unique to OHC vs. O such as ECM-receptor interaction, Hippo signalling, and PI3K-AKT signalling were predominantly located within the OHC-like cell cluster. Conversely, pro-inflammatory pathways including CLR pathway, osteoclast differentiation, and lipid and atherosclerosis that were uniquely augmented in OC vs. O were distributed around the OC-like cell cluster located within the main body of osteoclasts with little overlap with OHC-like cells. To further validate the identity of OHC-like cluster as our hybrid-like cells, we highlighted significant gene functions and pathways that coincide with the transcriptome of our hybrid-like cells (Fig. 6e). We also quantitatively scored the GSVA pathway profile for the OHC-like cluster against OHC and OC profiles separately to show that this cluster is significantly more similar to hybrid-like cells (Fig. 6f). These findings validated the presence of osteoclast-tumor hybrid-like cells *in vivo* in breast cancer patients with bone metastases.

### Hippo “off” pathway in osteoclast-tumor hybrid-like cells

To further explore the influence of pathways enriched in hybrid-like cells, we pinpointed main transcription factor (TF) drivers responsible for activating these pathways in OHC and OC (Fig. 7a). While both cell types shared several TFs linked to canonical osteoclastogenic and stress-response programs such as Jun, Hif1a, and Nfkb1, the hybrid-like cells exhibited a distinctive signature marked by strong Tead1 activity and Yap1/Taz expression (data not shown). These gene signatures imply Hippo pathway inactivation, which may alter osteoclast proliferation, motility, and ECM remodeling. Using the same co-culture experiment described in Fig. 5, we isolated Cd45+ osteoclasts and analysed their alterations on osteoclast enzymes, ECM components, and Hippo “off” genes under direct contact versus paracrine conditions (Fig. 7b). While no alterations were observed for osteoclast enzymes, we showed that expression levels for ECM components (Fn1, Pcolce2, Col4a2) and Hippo inactivation factors (Yap1, Amolt2, Ankrd1, Areg, Ctgf, Cyr6l, Itga2, Serpine2) were markedly activated to a higher level under direct contact with tumors. Notably, Cd45+ osteoclasts from direct contact consisted of a mixture of OHC- and OC-like cells. Our Smart-Seq 2 data analysis suggests that these ECM component and Hippo inactivation were likely induced by the presence of hybrid-like cells.

**Fig. 7:**
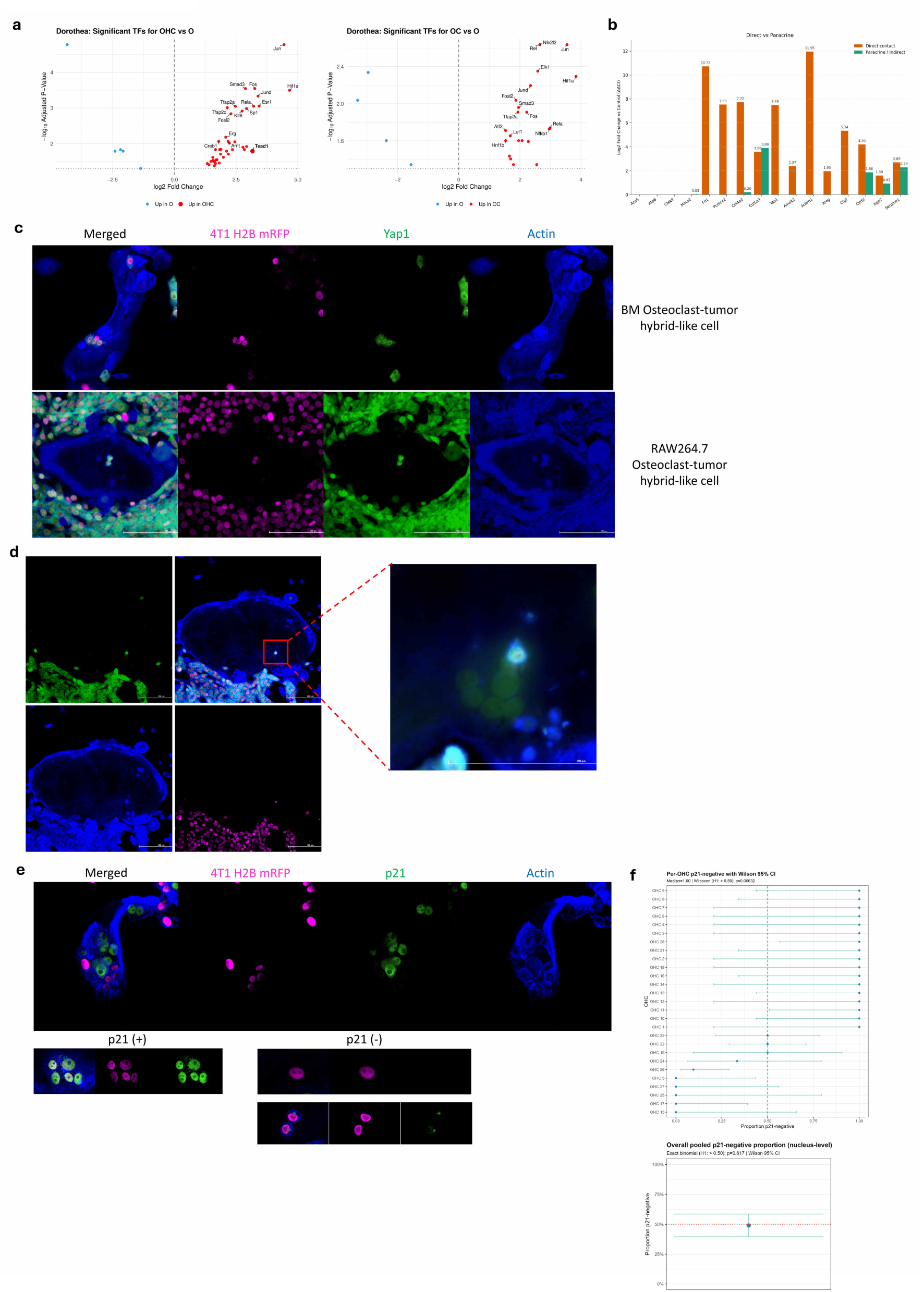
Yap1-activated osteoclasts exhibit a proliferation-permissive state. **a,** Dorothea analyses for OHC vs. O and OC vs. O highlighted Yap1-dependent TEAD1 transcription factor expression in hybrid-like cells. **b,** RAW264.7 osteoclasts from direct co-culture with 4T1 breast cancer cells promoted Yap1 downstream gene activation (Amotl2, Ankrd1, Ctgf, Cyr6I, Itga2, Serpine1) and enhanced ECM-modifying factors (Fn1, Pcolce2, Col4a2, Col5a3). **c,** Tumor nuclei present in primary and RAW264.7 hybrid-like cells displayed nuclear colocalized Yap1. **d,** Hybrid-like cell containing tumor nucleus with colocalized Yap1 were located adjacent to osteoclast nuclei expressing low Yap1 signals. **e,** Hybrid-like cells stained for p21 cell cycle inhibitor with p21-colocalized nuclei indicated as p21-positive (+) and non-p21-colocalized nuclei indicated as p21-negative (-). **f,** OHC-level quantification of p21-negative tumor nuclei, plotted as the fraction of p21-negative tumor nuclei per individual OHC with Wilson 95% confidence intervals; significance versus 50% was assessed by one-sample Wilcoxon signed-rank test (p = 0.00632). **g,** Nucleus-level quantification of p21-negative tumor nuclei pooled across OHCs, plotted as the overall p21-negative fraction with Wilson 95% confidence intervals; significance versus 50% was assessed by exact binomial test (p = 0.617).

To validate whether hybrid-like cells truly possess Yap1 in its activated form, we assessed Yap1 by staining for its presence in direct co-cultures from primary BM and RAW264.7 osteoclasts against the appropriate breast tumors (Fig. 7c). We detected strong Yap1 signals that colocalized with the majority of tumor nuclei found within hybrid-like cells and in individual tumor cells. Comparatively, we observed most non-tumor nucleus (negative for H2B-mRFP) in osteoclasts and hybrid-like cells having very dim to no Yap1 presence. This observation suggests that Yap1 expression in hybrid-like cells originated from tumor nuclei themselves. To infer how this may potentially influence transcriptional machinery in hybrid-like cells, we showed that these Yap1+ tumor nuclei seemingly integrate adjacent to the host osteoclast nuclei (H2B-mRFP-) with low Yap1 signals (Fig. 7d). Concurrent with our transcriptome findings, the inclusion of tumor nuclei in hybrid-like cells may alter their cellular functions and phenotype. As Yap1 activation drives proliferation and have an inverse correlation with p21 cell cycle inhibitor expression, we stained and quantified for p21 expression in tumor nuclei from hybrid-like cells (Fig. 7e). We identified tumor nuclei with strong p21 colocalization signals as p21+ tumor nuclei and signals without nucleus colocalization as p21- tumor nuclei. At the OHC level where we treated each hybrid-like cell as a single unit and scored by the fraction of its tumor nuclei that were p21-negative, the proportion of p21- tumor nuclei were significantly greater than 50% (p = 0.00632), indicating that a typical OHC contains predominantly p21-negative tumor nuclei. In parallel, we also quantified p21 absence at the tumor nucleus level by pooling all tumor nuclei across OHCs to determine the overall fraction of p21- nuclei. This complementary analysis was performed to distinguish whether p21 negativity was enriched on a per-hybrid-cell basis or predominated across the entire nucleus population. Under the pooled nucleus-level view, 50 of 102 tumor nuclei (49.0%) were p21-and this was not significantly different from 50% (p = 0.617) (Fig. 7f). Together, these data indicate that individual OHCs are frequently enriched for p21- tumor nuclei, but not at the pooled nuclei level. These findings are consistent with hybrid-like cells existing in a more proliferation-permissive state than non-hybrid osteoclasts. However, whether and how the incorporation of p21-negative tumor nuclei functionally contributes to the mitotic machinery of hybrid-like cells remains unclear.

## Discussion

The role of osteoclasts and cells of myeloid lineage in bone metastasis has mostly been documented on paracrine interactions via cytokines and other factors contributing to vicious tumor growth and osteolysis[21]. Recent spotlight on cell fusion between cells of myeloid lineage and tumor cells suggests that physical cell-cell interactions may contribute to outcomes in metastatic progression[1, 22, 23]. In our study, we showed that solid tumors can merge with osteoclasts in various differentiation stages via dynamic cell fusion and cell-in-cell mechanics, generating a hybrid-like cell type that possesses unique properties that can potentially have profound effects on bone metastasis. We further showed that tumor nuclei in our hybrids were transcriptionally active. A similar phenomenon was described by Andersen et. al., who reported that osteoclasts from myeloma (a blood cancer of B cell origin) patients with bone metastases contained transcriptionally active tumor nuclei formed by cell fusion[10]. Our data support that osteoclast-tumor fusion is not restricted to blood cancers. On this note, osteoclasts that harbor tumor cells in their cytoplasm have been described in thyroid cancer[9]. Although cell-in-cell models have been lacking in osteoclast studies, we surmise that osteoclasts containing solid tumors within their cytoplasm are formed through dynamic combination of cell fusion and cell-in-cell phenomenon. Further in-depth cellular dynamics studies are required to fully comprehend how both processes were regulated to form hybrid-like cells.

Osteoclasts exhibit diversified functions under inflammatory and disease conditions[24]. Our gene expression profiling classified osteoclast-tumor hybrid-like cells as an osteoclast variant that has enhanced pro-angiogenic and ECM-remodelling functions. Angiogenesis is important for tumor vasculature to acquire nutrient and oxygen support[25], while deregulated ECM components lead to disorganized bone matrices for tumor invasion[26]. Together, these two alterations can induce mechanotransduction changes on tumor cells that promotes tumor survival via matrix stiffness[27]. We propose that hybrids may directly modulate tumor colonization in bone metastases. In addition to functional changes, our study showed that hybrid-like cells express cancer associated PI3K-AKT and Hippo effectors. While PI3K-AKT signalling is notorious for its cancer-promoting functions, it is also essential for osteoclast survival[28]. On the other hand, Hippo effectors (YAP1/TAZ-TEAD) activate transcription programs involved in proliferation, EMT, and chemoresistance[29]. These pathway activity patterns may confer survival advantages in hybrid-like cells over non-hybrid osteoclasts. Consistent with this, the enrichment of p21- tumor nuclei in hybrid-like cells suggests a more proliferation-permissive state. However, our study cannot determine whether these hybrids are oncogenic.

We found that breast cancer cells were influenced by the presence of osteoclasts to express hematopoietic lineage genes in relation to osteoclast/lymphocyte differentiation and antigen presentation functions. Tumor cell transdifferentiation into other cell lineages has been described in various cancers. Glioblastoma has been shown to differentiate into neuron-like cells, and prostate cancer is known to undergo osteomimicry to become osteoblast-like cells[30]. Furthermore, various cancers exhibit vascular mimicry by differentiating into endothelial-like cells[31, 32]. In our study, breast cancer cells required osteoclasts to be present for differentiation to occur. Remarkably, CTCs in patients with bone metastases have been reported to express osteoclast differentiation genes[33]. We theorize that changes in these CTCs may be attributed to their prior interactions with osteoclasts and/or their precursors. Collectively, we propose that breast cancer cells undergo a process similar to osteomimicry, which alters their morphologies from adherent, epithelial cells into detached, round cells prior to dynamic fusion/CIC entry into osteoclasts. This shift to round morphology is also observed in osteoclast fusion[34].

Our results demonstrated that physical contact between tumors and osteoclasts are not only necessary for cell fusion and CIC processes, but also to modify osteoclast functions. This is in line with Gu et al.’s observations, which found that tumor cells can transfer cellular material to osteoclast precursors through migrasomes to initiate osteoclast differentiation[13]. We acknowledge that tumor cells in our mice study may also acquire Cre activity through similar mechanisms. Nonetheless, tumor cells likely require close proximity to osteoclasts for either cell fusion or migrasome-mediated transfer to undergo Cre-Lox recombination to express GFP.

Beyond enabling physical exchange processes, direct tumor–osteoclast contact also appeared to reshape osteoclast behavior. Interestingly, hybrid-like cells and osteoclasts in direct contact with tumor cells appeared to shift away from a predominantly osteolytic state toward a distinct tumor-educated phenotype. Although breast tumors are known to exarcebate osteolysis by activating osteoclasts[35], our data suggests that osteoclasts exposed to direct cell–cell contact and fusion/CIC with breast tumor cells preferentially adopt ECM-remodelling and immunoregulatory features, reminiscent of the phenotypic reprogramming observed in tumor-associated macrophages (TAMs). This interpretation is consistent with the macrophage lineage origin of osteoclasts.

In conclusion, our study unveiled that breast tumors participate in osteoclast fusion and/or CIC processes to generate osteoclast-tumor hybrid-like cells. These hybrid-like cells exhibit immunoregulatory, pro-angiogenic, and ECM-remodelling functions and express cancer-associated PI3K-AKT and Hippo pathways. In bone metastases, this mechanism provides additional immune evasion pathways where breast tumors can camouflage themselves to resemble osteoclasts and by modifying osteoclast phenotype in a similar manner to TAMs. Our findings suggest that these hybrid-like cells may contribute to the establishment and progression of metastatic bone lesions, highlighting their potential for therapeutic interventions.

## Methodology

### Cell Culture

RAW264.7, 4T1, EO771, B16F10, and CT26 cell lines were obtained from ATCC. Cells were cultured in RPMI 1640 medium supplemented with 10% FBS and 1% penicillin/streptomycin at 37°C with 5% CO_2_. Primary bone marrow (BM) cells used for osteoclast differentiation were isolated from C57BL/6 mice. Bone marrow derived macrophages (BMMs) were differentiated from BM cells using 25 ng/ml mM-CSF. On day 3, BMM medium was replaced with 10 ng/ml mRANKL, 25 ng/ml mM-CSF, and 40 ng/ml mTNF-α and cultured for 3 more days to differentiate BMMs into osteoclasts. RAW264.7 macrophages were differentiated into osteoclasts using 100 ng/ml mRANKL, 75 ng/ml mM-CSF, and 40 ng/ml mTNF-α for 5 days, with medium replaced on day 2.

### Lentiviral transduction

RAW264.7 macrophages were transduced with lentiviral particles generated from pLenti-CMV-GFP-Hygro vector to produce RAW264.7-GFP cells after hygromycin selection. 4T1, EO771, B16F10, and CT26 were transduced with lentiviral particles generated from pHIV-H2B-mRFP to produce cancer cell lines with H2B-mRFP. Cancer cell lines with H2B-mRFP were sorted using a Sony SH800Z cell sorter. Cre-reporter cell lines were generated by transducing EO771 and B16F10 with lentiviral particles harbouring a Cre reporter (floxed DsRed-E2 with downstream eGFP) followed by puromycin selection.

All plasmids were purchased from Addgene: pLenti-CMV-GFP-Hygro (656-4) (Addgene #17446), pHIV-H2BmRFP (Addgene #18982), Cre Reporter (Addgene #62732).

### Osteoclast precursor-cancer cells co-cultures

RAW264.7-GFP and 4T1-H2B-mRFP cells were seeded at a 20:1 ratio on day 0. mRANK-L (100 ng/ml), mM-CSF (75 ng/ml), and mTNF-α (40 ng/ml) were added in the culture media on the same day. Medium was replaced using the same cytokines on day 2. Cells were co-cultured for up to day 5. For co-cultures with other cancer cell lines, cell seeding density was adjusted according to their respective doubling time.

Differentiated primary BMMs were mixed with EO771 Cre reporter at a 37:1 ratio. Cytokine composition used in primary osteoclast differentiation assay was added into the culture media on the same day. Cells were co-cultured up to day 3.

Co-cultured RAW264.7 and 4T1 cell isolation for downstream assays were magnetically sorted twice using CD45 MicroBeads, mouse (Miltenyi, 130-052-301) to acquire pure RAW264.7 osteoclasts and 4T1 breast cancer cells as CD45+ and CD45- cell fractions respectively.

### Western Blot

Proteins were extracted from cells, quantified via BCA assay, separated by SDS–PAGE, and transferred to membranes by wet transfer. Membranes were blocked using 1% BSA and probed with anti-PU.1/Spi1 (31 kDa) (rabbit recombinant monoclonal, clone EPR22624-20, Abcam, ab227835) at 1:1000 dilution and anti-β-actin (42 kDa) (mouse monoclonal, clone AC-15, Sigma-Aldrich, A5441) as a loading control at 1:5000 dilution. Detection was performed using species-appropriate HRP-conjugated secondary antibodies and Clarity Western ECL Substrate (Bio-Rad, #1705061) according to the manufacturer’s instructions.

### RT-qPCR

RNA was isolated using FastPure Cell/Tissue Total RNA Isolation Kit V2 (#RC112) according to the manufacturer’s instruction. Collected RNA was transcribed into cDNA using the PrimeScript RT Master Mix Kit (Takara Bio, Inc.). Quantitative PCR (qPCR) was performed using PowerUp™ SYBR™ Green Mix (Applied Biosystems™, A25742). Beta-actin was used as the reference gene for normalization, and relative changes in gene expression were determined using 2−ΔΔCt method. Primer sequences used for qPCR are shown in Supp. Table 1.

### Immunocytochemistry

Cells were fixed with 4% PFA solution and permeabilized using 0.1% Triton-X solution. Fixed and permeabilized cells were blocked using 1% BSA and 300 uM glycine in PBS with 0.1% Tween-20 for 30 minutes. Cells were then subjected to primary antibody staining overnight at 4°C, followed by staining with secondary antibodies for 1 hour at room temperature. Antibodies: anti-RFP (Rockland 200-101-379), anti-H3K4me2 (Invitrogen PA5-114685), anti-GFP (Aveslabs GFP-1020), Yap (D8H1X) XP Rabbit mAB (CST-14074S), Anti-p21 antibody (ab188224), AlexaFluor488 AffiniPure™ donkey anti-rabbit IgG (H+L) (Jackson Immunoresearch, 711-545-152), AlexaFluor Plus 647 donkey anti-goat IgG (H+L) (Invitrogen A32849), and AlexaFluor Plus 405 Phalloidin (Invitrogen A30104), Alexa Fluor 488 AffiniPure donkey anti-chicken IgY (IgG) (H+L) (Jackson Immunoresearch 703-545-155). Cells were imaged on a Nikon A1HD25 Confocal Microscope with Nikon NIS Elements for data collection. Images were further processed using ImageJ Fiji.

### Time-lapse live cell recording

The CytoSMART™ Lux3 FL fluorescence live cell imager was used to monitor cell fusion in co-culture. Cells were recorded through brightfield and green/red fluorescence imaging from day 0-5 with adjustments to track cells of interest. Snapshot images and time-lapse recording were automatically processed via Axion Biosystems online server system.

### RAW264.7-GFP-4T1-H2B-mRFP co-culture on laser microdissection dish

Cells were co-cultured on specialized dish made with polyethylene naphthalate (PEN) membrane designed for laser microdissection. PEN Petri Dishes 4 µm (11600295) were obtained from Leica. Cells were seeded and spread evenly across the culture area. Growth medium supplemented with osteoclastogenic cytokines was added drop-by-drop to minimize disruption on cell distribution. To prevent dehydration, dishes were placed in a 10 cm^2^ dish with the peripheral filled with PBS. Medium was replaced on day 2 and day 4. On day 5, cells were used for laser microdissection process.

### Laser microdissection

Leica microdissection microscope (LMD 7000) was used to isolate osteoclast-tumor hybrid-like cells and other cells of interest. Cells were identified based on fluorescence prior to excision. Osteoclast-tumor hybrid-like cells were recognized as GFP+ osteoclasts with intact mRFP+ tumor nuclei; non-hybrid osteoclasts were recognized as GFP+ osteoclasts without signs of mRFP+ tumor cells; tumor cells were recognized as mRFP+ single cells. Excised cells were immediately collected in lysis buffer containing 2.5 mM dNTP mix (Invitrogen) (R0192), 1 U/µl SUPERase·In™ RNase Inhibitor (Invitrogen) (AM2694), and 0.1% Triton™ X-100 (Sigma) (9036-19-5) in RNAse-free ddH_2_O. Cells were then immediately frozen in dry ice and stored at -80°C.

### Cytokine array

Proteome Profiler Mouse XL Cytokine Array (Biotechne) was used to assess cytokine/chemokine profile from supernatant collected from co-culture according to manufacturer’s instructions. The membrane images were visualized the ChemiDoc Imaging system (Bio-Rad), and the intensity of each spot in the membranes was analyzed with Image Lab software (Bio-Rad).

### Mouse experiment

All mouse experiments were approved by the Animal Experimentation Ethics Committee of the City University of Hong Kong (Ref. No.: AN-STA-00000207). Mice were housed in a laboratory environment at 24 ± 1 °C under a 12 h dark-light cycle and fed a standard chow. Ctsk-2A-Cre mice (Cat. NO. NM-KI-190019) were purchased from Shanghai Model Organisms Center, Inc. WT C57BL/6 mice was provided by Laboratory Animal Research Unit of City University of Hong Kong (ISO 9001 verified). Mice of 8-12 weeks old were used for the bone metastasis model. Male and female mice were used for B16F10 and EO771 bone metastasis model respectively. Genotyping for Ctsk-Cre mice was performed using the following primers: *Ctsk-cre (Mut)* (GAGAGCTGGGGAAACAAAG, GGTAGTCCCTCACATCCTCAG); *Ctsk-cre (Wt)* (TCTACCTTCAAAGTGCTGCCATTA, AGAGAAGGGAAGTAGAGTTGTCAC).

### Bone metastasis model via caudal artery injection

Bone metastasis was induced as described by Kuchimaru, T., et al.[36] Briefly, mice were sedated with pentobarbital sodium via intraperitoneal injection. 250,000 cells in 100 µl volume were injected through the caudal artery at the tail. Mice were monitored for bone metastatic development as determined by an in vivo imaging system (IVIS® Lumina Series III). Prior to imaging, mice were shaved to eliminate autofluorescence. DsRed fluorescence in EO771 and B16F10 Cre reporter was detected. Background and autofluorescence were removed via spectral unmixing fluorescence. Biological replicates of n≥10 was used for each tumor group.

### Tumor and bone marrow processing in mice

Mice were sacrificed upon signs of suffering or pain. Bone metastases were collected and dissociated into single cells using Singleron PythoN® Tissue Dissociation System. BM cells from hind limbs with bone metastases were flushed and collected as single cell suspension. Biological replicates of n≥4 was used for bone metastasis samples while biological replicates of n≥3 was used for bone marrow samples.

### Flow cytometry

Cells were blocked in TruStain FcX™ (anti-mouse CD16/32) Antibody (Biolegend) (101320) to prevent non-specific bindings. Live cells were assessed directly for DsRed and GFP fluorescence. For fixed cells, BD Cytofix/Cytoperm™ Fixation/Permeabilization Kit (BD Biosciences) (554714) was used prior to staining for GFP and DsRed fluorescent proteins. Anti-RFP (Rockland 200-101-379) was used to stain DsRed fluorescent protein. Flow cytometry data were expressed as mean ± SD. Statistical significance was evaluated by a two-tailed unpaired Mann-Whitney U test unless otherwise indicated. P values <0.05 were considered statistically significant. Figures were made with GraphPad Prism software.

### SMART-seq 2 single cell transcriptome sequencing

Frozen single cell RNA samples were submitted to BGI Genomics for sample quality control and SMART-Seq 2 sequencing. Each cell type had n ≥ 5 biological replicates. Sequencing was performed using high throughput DNBSEQ platform with a PE100 sequencing strategy for full transcriptome analysis.

### RNA sequencing data preprocessing

Raw reads were first quality controlled using FastQC[37]. The validated reads were then aligned to the Mus musculus genome (genome-build-accession NCBI:GCA_000001635.8) using STAR[38] with default parameters. RSEM[39] was used to quantify the gene expression level. PCA was performed to evaluate the batch effect and to visualize the sample distribution. Differentially expressed genes (DEGs) were detected for each comparisons using DESeq2[40]. Gene Ontology and KEGG pathway enrichment analyses were performed using clusterProfiler [41]. The R package ggplot2 was used to visualize the results[42].

### Public Database Comparison

Single cell sequencing data from a patient with ER+ breast cancer bone metastasis (GSE190772) was used[20]. Osteoclast and epithelial tumor populations were extracted from the database. The extracted data was normalized through the Seurat package (v5.1.0)[43], and a UMAP was created for each cell population. Gene annotations from Homo sapiens were matched with *Mus musculus* for further analysis.

Gene set variation analysis (GSVA; v1.52.3)[44] was conducted to compare DEGs in our data to individual cells in osteoclast and epithelial populations from bone metastasis respectively. Additional GSVA was performed on osteoclast and epithelial tumor population for specific KEGG pathways of interest with DEGs from our data.

## Funding

This work was supported by the following funding sources: Croucher Foundation (Ref. No. 9509002 and 9500032)

The Science and Technology Innovation Committee of Shenzhen Municipality (Ref. No. JCYJ20210324133813036)

Hong Kong Research Grants Council (Ref. No. C7008-22G and 11103523) Tung’s Biomedical Sciences Centre (Ref. No. 9609306)

City University of Hong Kong (Ref. No. 7005313, 7005518, 7005740, and 7005873)

## Author contributions

Conceptualization: K.H.L. and K.T.C. Methodology: K.H.L., D.S., and X.L. Investigation: K.H.L., D.S., X.L., E.D., M.W., and Y.L. Visualization: K.H.L., D.S., and X.L. Supervision: K.T.C. and X.W. Writing: K.H.L. and K.T.C.

## Competing interests

All authors declare they have no competing interests.

## Data and materials availability

All data are available in the main text or the supplementary materials.

---

**Fig. S1.**
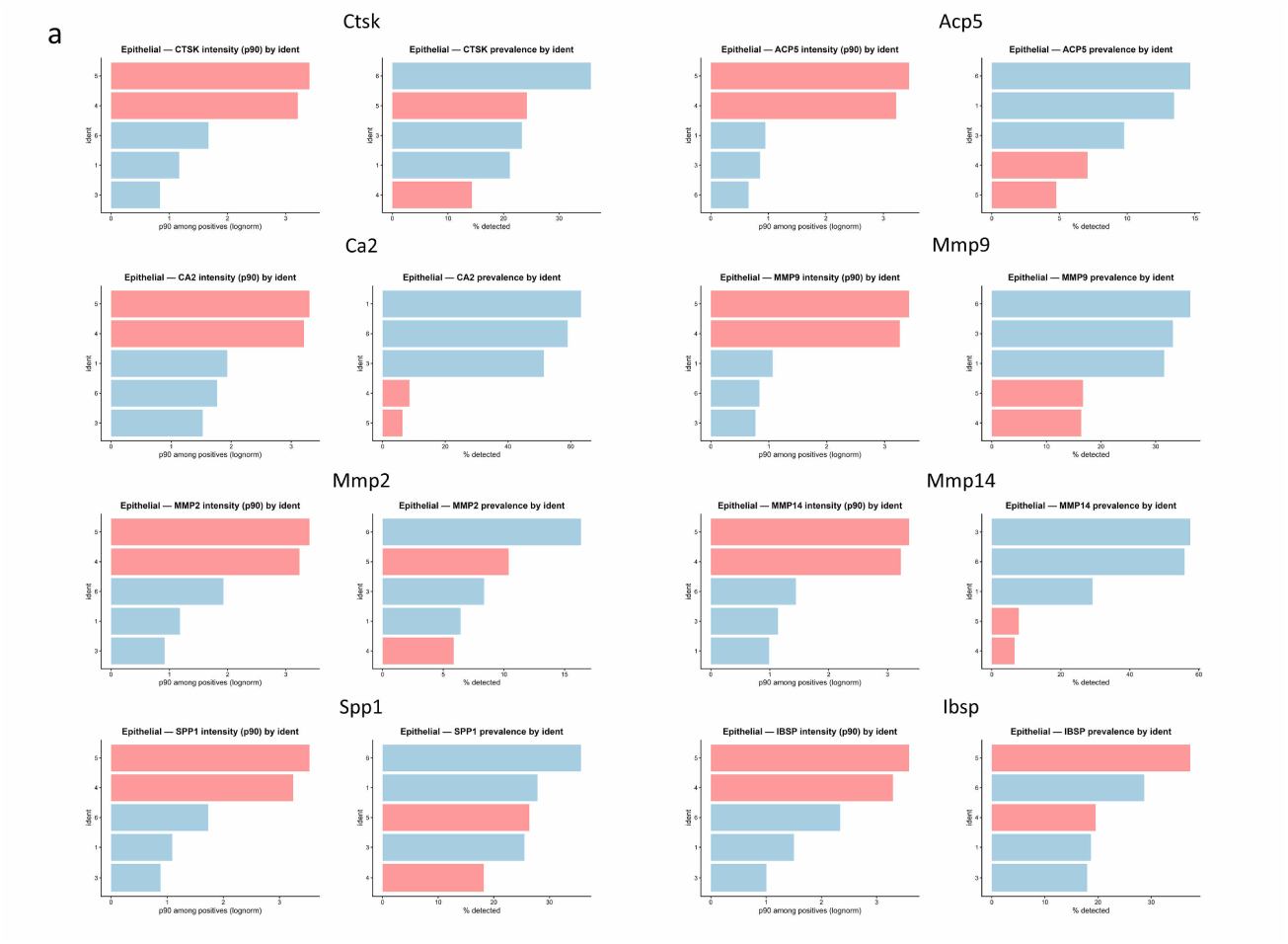
OC/OM gene-set prevalence and intensity across epithelial subclusters. a, Prevalence (% detected) and intensity (p90 among positive cells) of OC/OM-associated genes in epithelial subclusters 1, 3, 4, 5, and 6. Genes shown include CTSK, ACP5, CA2, MMP9, MMP2, MMP14, SPP1, and IBSP. Statistical significance was assessed using the Wilcoxon rank-sum test.

**Fig. S2.**
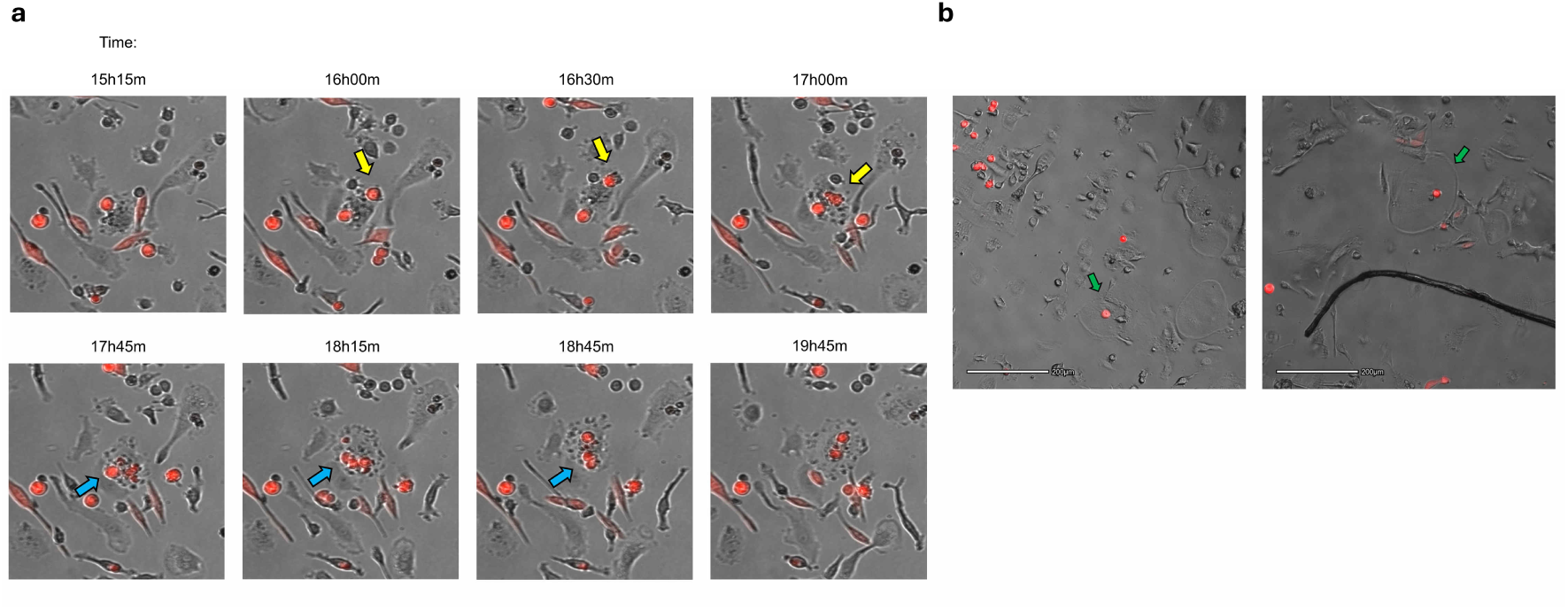
Primary osteoclasts fuse with breast cancer cells *in vitro*. a, Primary osteoclast acquiring EO771 Cre reporter and their nuclei into its cytoplasm (yellow & blue arrows). Video stills were acquired from Movie S4. b, Examples of EO771 Cre reporter tumor nuclei present inside developing primary osteoclasts (green arrows).

**Fig. S3.**
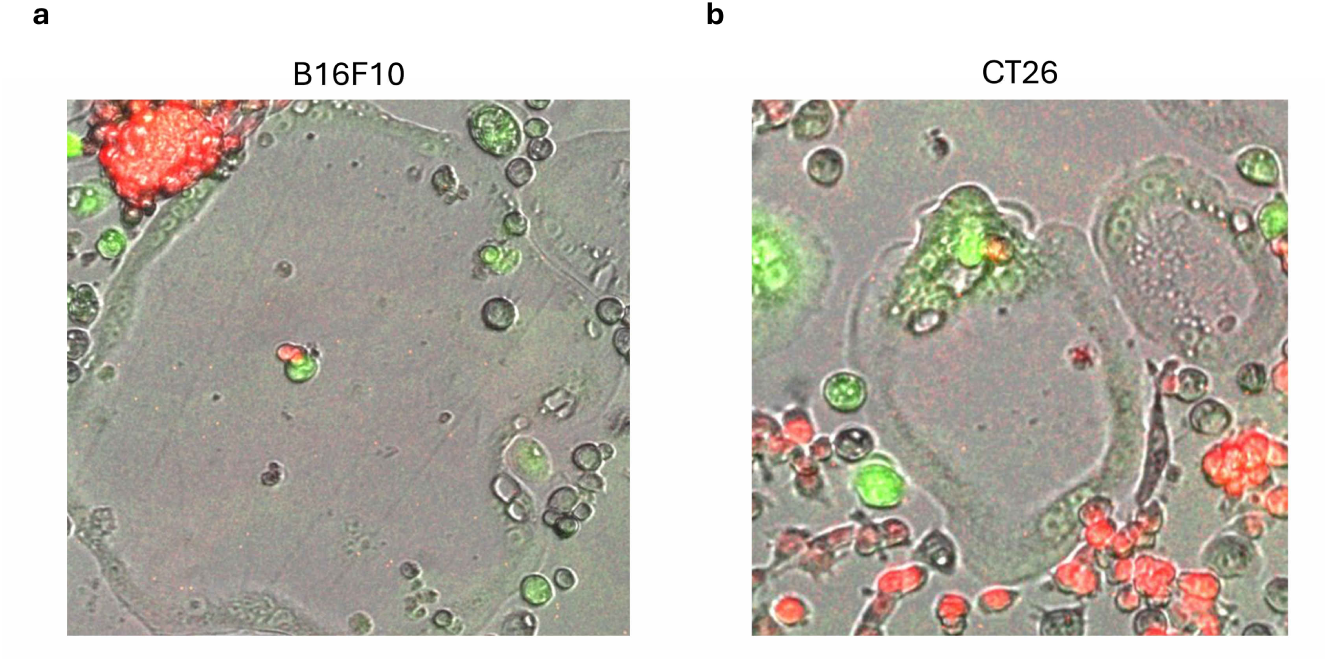
Osteoclasts fuse with melanoma and colon cancer cells *in vitro*. a,b, Fluorescent snapshot image for osteoclast-tumor hybrid-like cells formed between RAW264.7-GFP and other solid tumor cell lines. Hybrids containing tumor nuclei derived from (a) B16F10-H2B-mRFP melanoma and hybrids with tumor nuclei derived from (b) CT26-H2B-mRFP colon cancer.

**Fig. S4.**
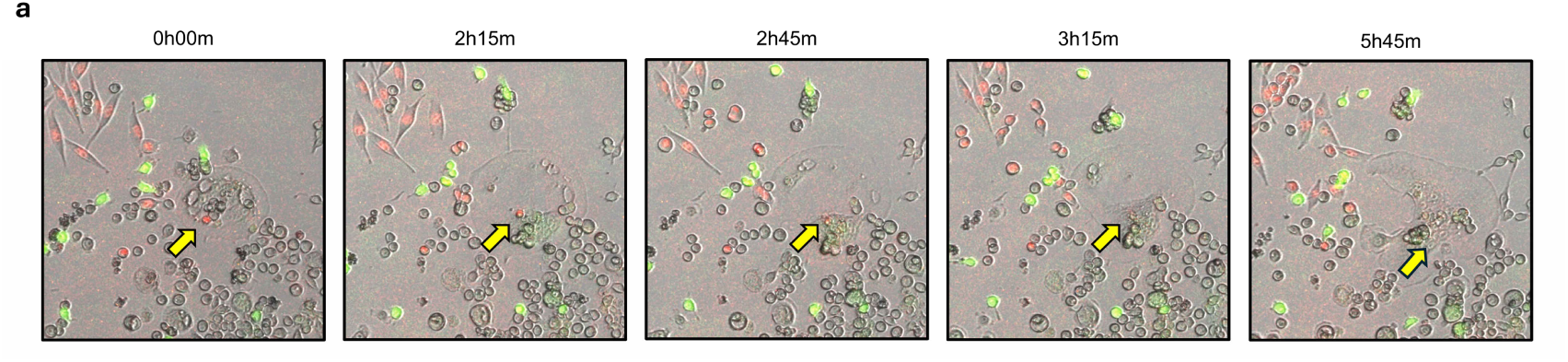
Cell-in-cell phenomenon between tumor and osteoclast. a, Tumor nucleus migrated near osteoclast nuclei and showed signs resembling disintegration with declining tumor fluorescent signal (yellow arrow).

**Fig. S5.**
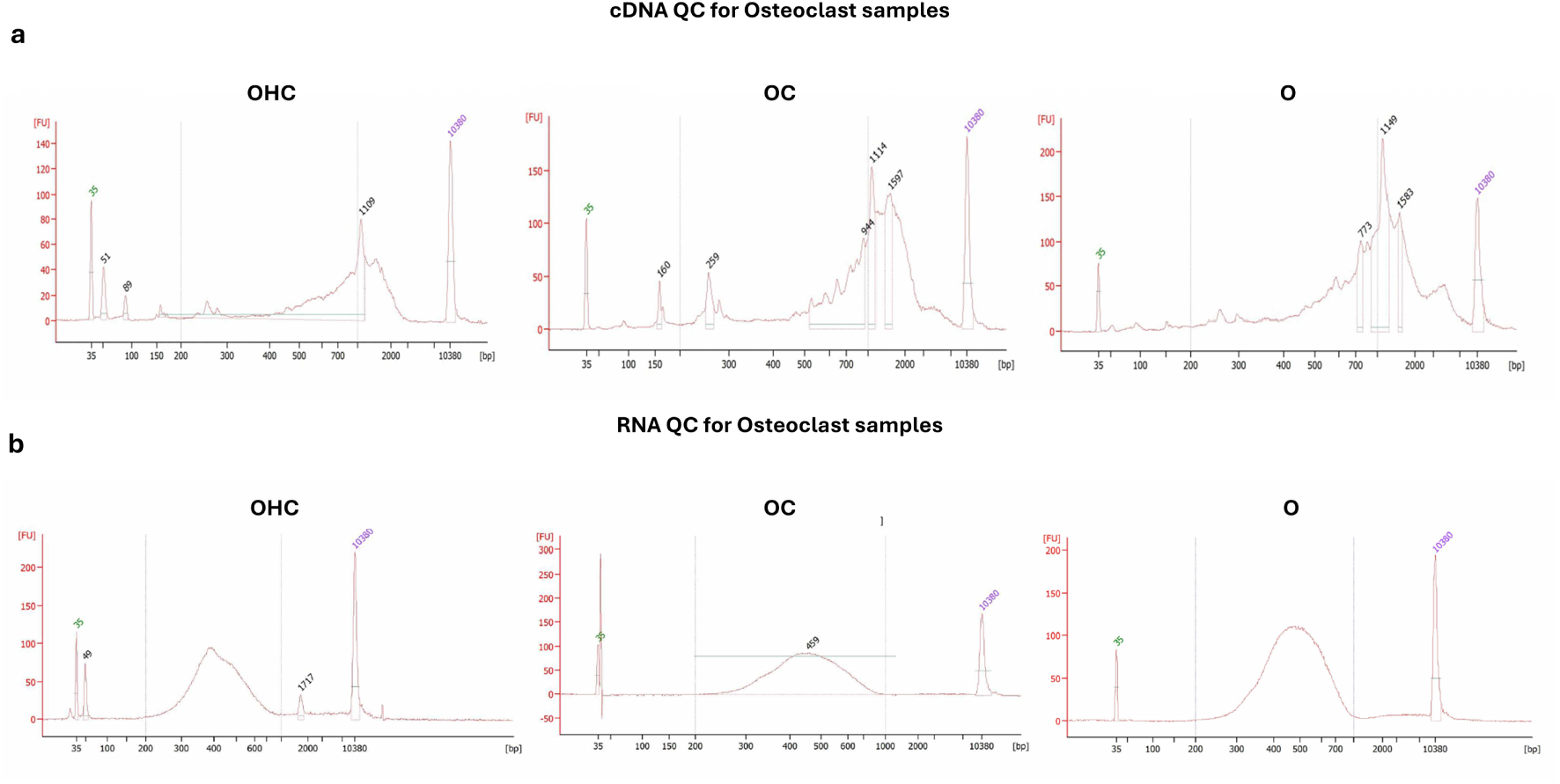
RNA quality control for osteoclast variants. a,b, cDNA and library QC graphs for three osteoclast sample types. (a) Representative cDNA QC graph for OHC, OC, and O. (b) Representative library QC graph for OHC, OC, and O.

**Fig. S6.**
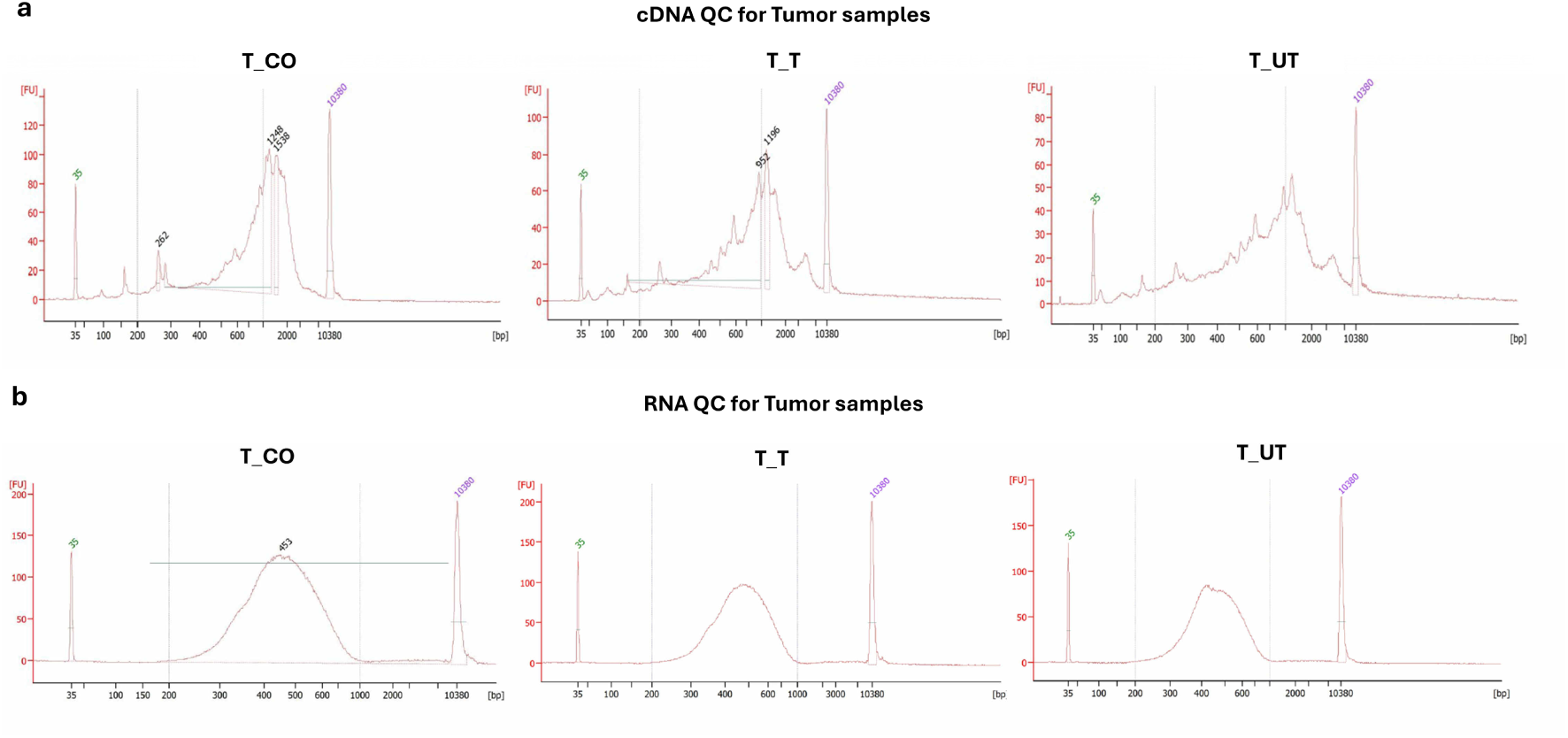
RNA quality control for tumor variants. a,b, cDNA and library QC graphs for three tumor sample types. (a) Representative cDNA QC graph for TUMOR_CO, TUMOR_T, and TUMOR_UT. (b) Representative library QC graph for TUMOR_CO, TUMOR_T, and TUMOR_UT.

**Fig. S7.**
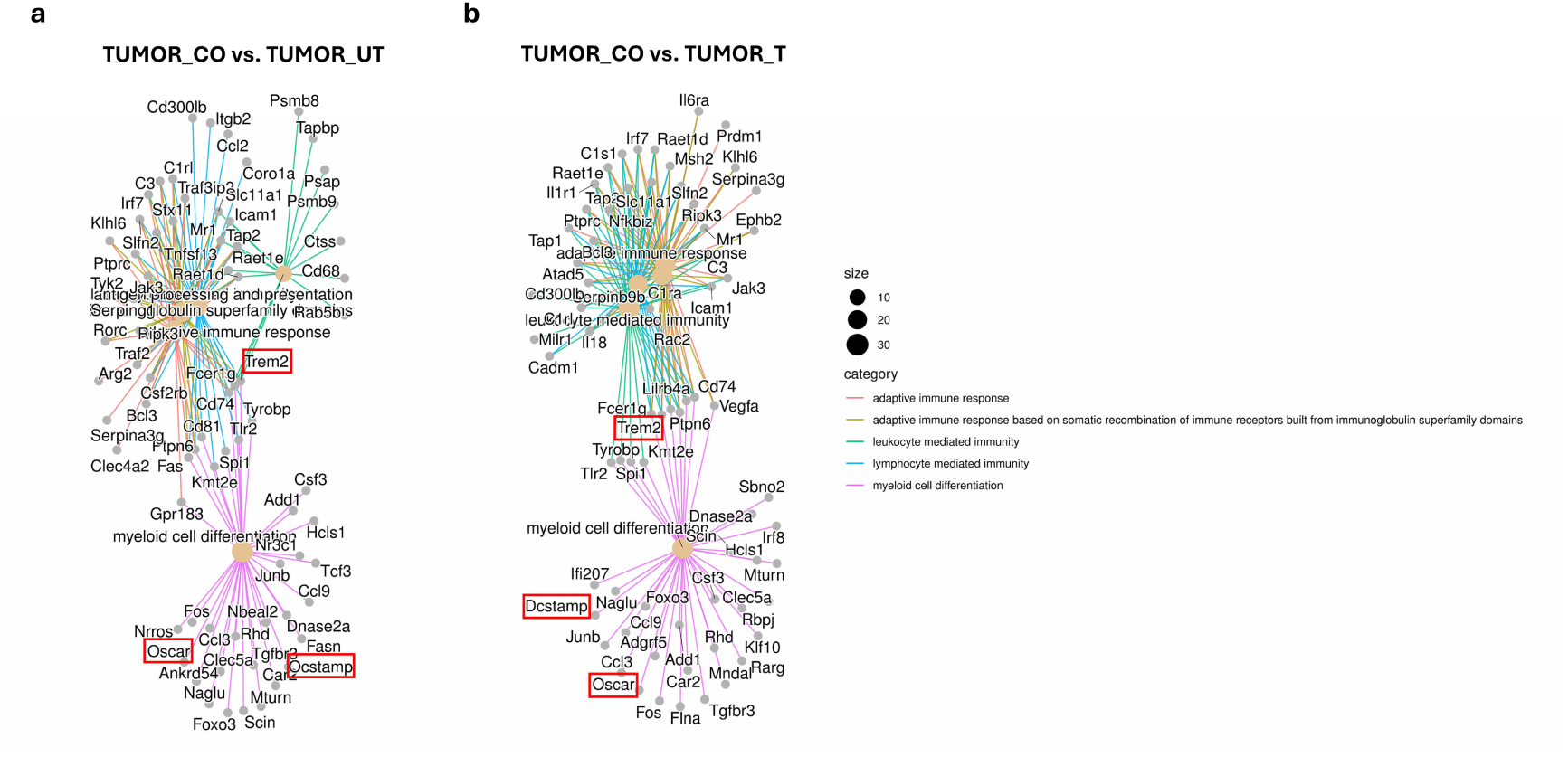
Gene ontology network analysis for DEGs in TUMOR_CO vs. TUMOR_UT and TUMOR_CO vs. TUMOR_T. a,b, GO network for DEGs from (a) TUMOR_CO vs. TUMOR_UT and (b) TUMOR_CO vs. TUMOR_T under different categories. Notable genes relevant to osteoclast differentiation and fusion were highlighted in red boxes.

**Fig. S8.**
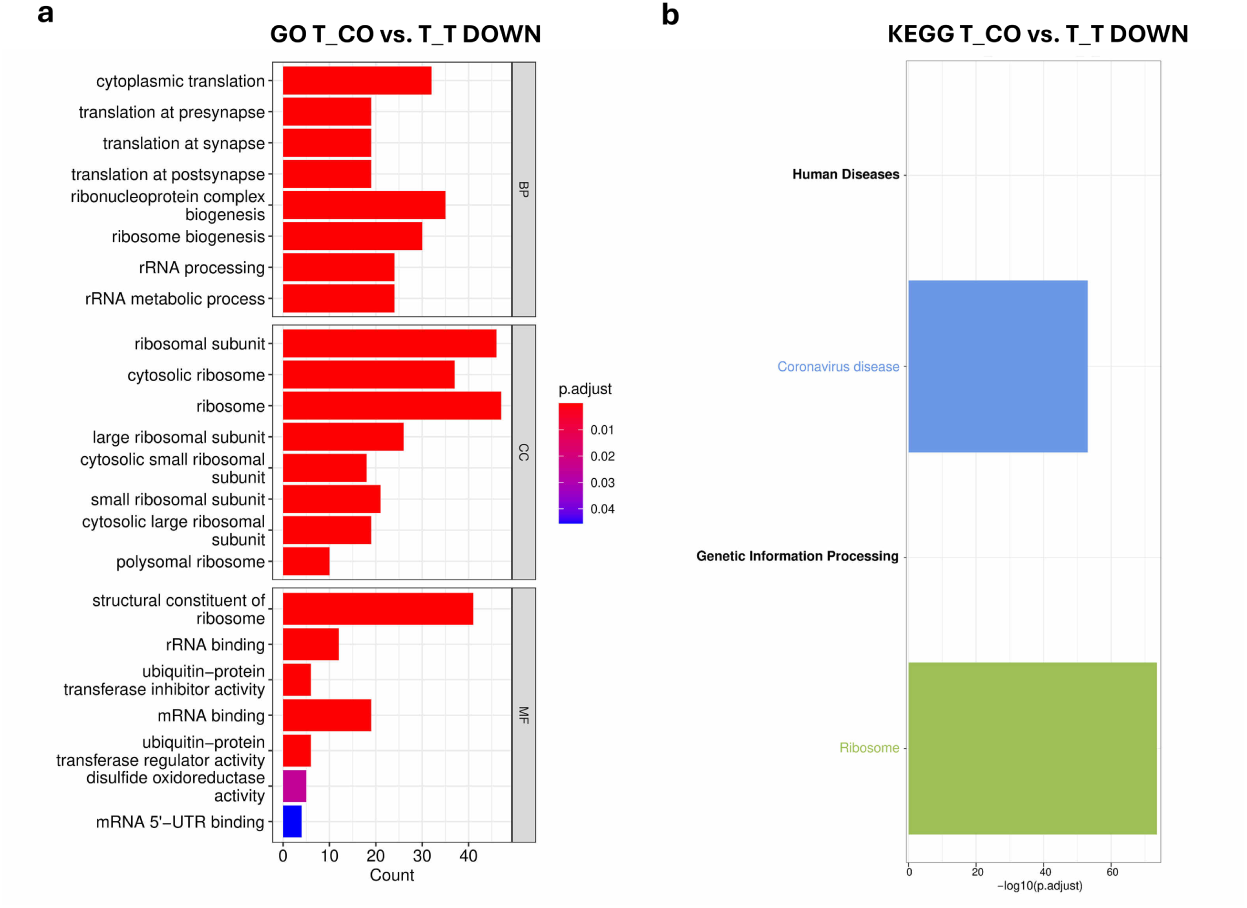
Attenuated genes and pathways in TUMOR_CO vs. TUMOR_T. a, GO analysis for downregulated DEGs in TUMOR_CO vs. TUMOR_T. GO categories shown included BP, CC, and MF. p.adjusted value < 0.05. b, KEGG pathway analysis for attenuated pathways in TUMOR_CO vs. TUMOR_T. p.adjusted value < 0.05.

**Fig. S9.**
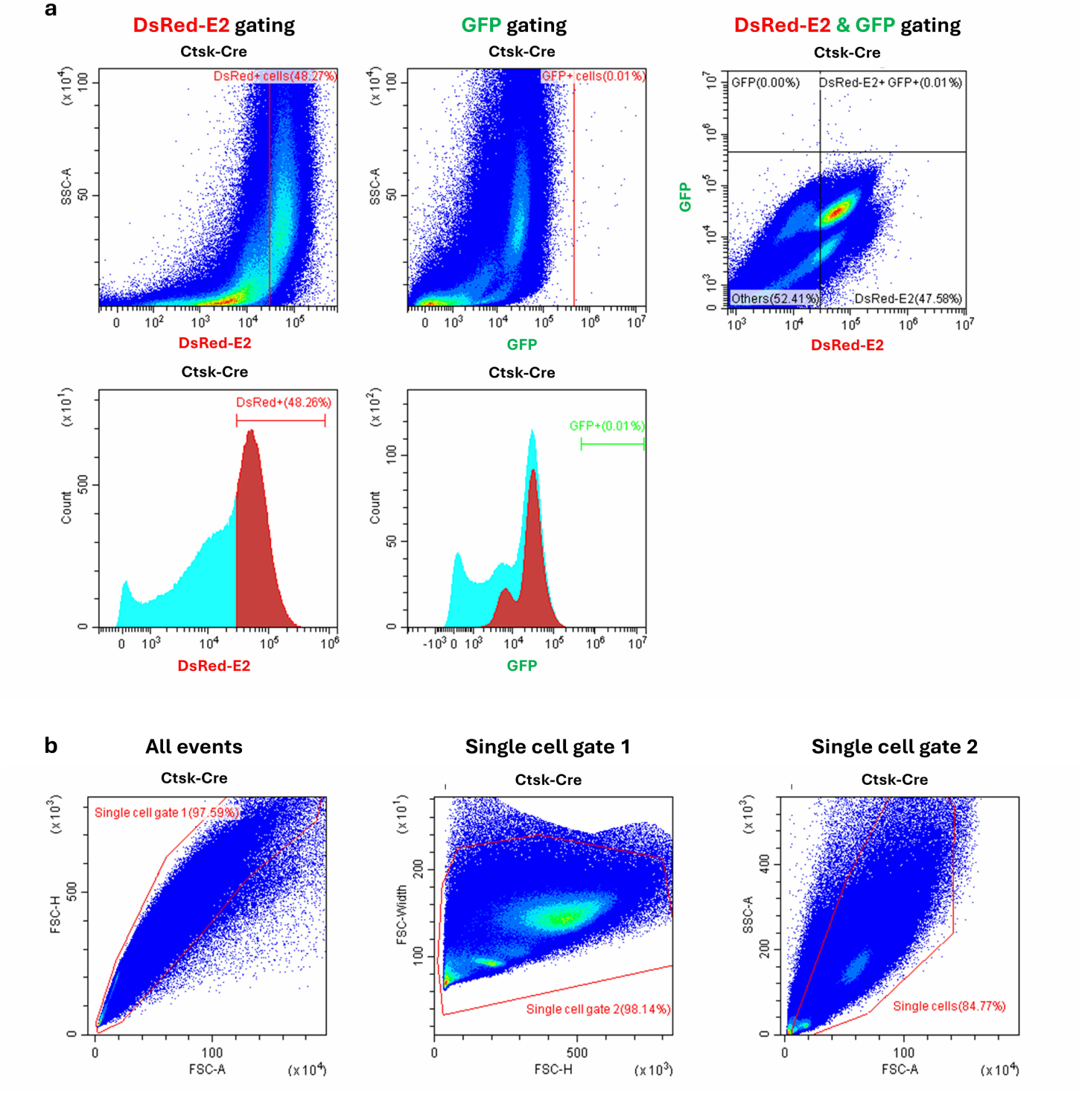
Flow cytometry analysis on bone metastases in Ctsk-Cre mice and single cell gating strategies. a, Flow cytometry analysis on gated single cells showing DsRed-E2+ and GFP+ cells in bone metastases derived from Ctsk-Cre mice. b, Single cell gating strategy on cells isolated from bone metastases. Doublet cells were gated out twice through FSC-A vs. FSC-H and FSC-H vs. FSC-W gates in Single cell gate 1 & 2 respectively. Cellular debris with low FSC-A & high SSC-A was gated out in Single cell gate 2 to acquire single cells for analysis.

**Movie S1. Live imaging of cell fusion between several pre-osteoclasts and an individual tumor cell.** 4T1-H2B-mRFP breast cancer cell fused and converged with RAW264.7-GFP pre-osteoclasts to form an osteoclast-tumor hybrid-like cell with intact tumor nucleus. Video was captured using CytoSMART Lux3 FL fluorescent time-lapse recording function. Video stills for Movie S1 were used in Fig. 1g.

**Movie S2. Live imaging of cell fusion between two osteoclast-tumor hybrids.** Two osteoclast-tumor hybrid-like cells fused with each other upon making cell membrane contact to form an enlarged hybrid-like cell. Video was captured using CytoSMART Lux3 FL fluorescent time-lapse recording function. Video stills for Movie S2 were used in Fig. 1h.

**Movie S3. Live imaging of individual tumor cells fusing into an existing osteoclast-tumor hybrid.** 4T1-H2B-mRFP breast cancer cells and RAW264.7-GFP fused and incorporated their nuclei into an existing osteoclast-tumor hybrid-like cell. Video was captured using CytoSMART Lux3 FL fluorescent time-lapse recording function. Video stills for Movie S3 were used in Fig. 1i.

**Movie S4. Live imaging of cell fusion between primary osteoclasts and breast cancer cells.** EO771 Cre reporter (red fluorescence) fused with primary BM-derived osteoclasts differentiated using BM cells from Ctsk-Cre mice. Video was captured using CytoSMART Lux3 FL fluorescent time-lapse recording function. Video stills for Movie S4 were used in Fig. .

**Movie S5. Live imaging of tumor nucleus activity in a hybrid-like cell post-fusion.** EO771-H2B-mRFP tumor nucleus disintegrated adjacent to osteoclast nuclei in an osteoclast-tumor hybrid-like cell. The hybrid proceeded to split into 3 daughter osteomorphs. Video was captured using CytoSMART Lux3 FL fluorescent time-lapse recording function. Video stills for Movie S5 were used in Fig. 2e.

## Notes

### Competing Interest Statement

The authors have declared no competing interest.

## References

1. Gast, C.E., et al., Cell fusion potentiates tumor heterogeneity and reveals circulating hybrid cells that correlate with stage and survival. Science Advances, 2018. 4(9): p. eaat7828.

2. Tretyakova, M.S., et al., Tumor Hybrid Cells: Nature and Biological Significance. Frontiers in Cell and Developmental Biology, 2022. 10.

3. Sieler, M., J. Weiler, and T. Dittmar, Cell-Cell Fusion and the Roads to Novel Properties of Tumor Hybrid Cells. Cells, 2021. 10(6).

4. Melzer, C., J. von der Ohe, and R. Hass, In Vivo Cell Fusion between Mesenchymal Stroma/Stem-Like Cells and Breast Cancer Cells. Cancers (Basel), 2019. 11(2).

5. Käkönen, S.-M. and G.R. Mundy, Mechanisms of osteolytic bone metastases in breast carcinoma. Cancer, 2003. 97(S3): p. 834–839.

6. Wang, M., et al., Molecular mechanisms and clinical management of cancer bone metastasis. Bone Research, 2020. 8(1): p. 30.

7. Tamura, T., et al., Extracellular Vesicles in Bone Metastasis: Key Players in the Tumor Microenvironment and Promising Therapeutic Targets. Int J Mol Sci, 2020. 21(18).

8. Gu, C., et al., Targeting initial tumour-osteoclast spatiotemporal interaction to prevent bone metastasis. Nat Nanotechnol, 2024.

9. Behzatoglu, K., Osteoclasts in Tumor Biology: Metastasis and Epithelial-Mesenchymal-Myeloid Transition. Pathology and Oncology Research, 2021. 27.

10. Andersen, T., et al., Osteoclast nuclei of myeloma patients show chromosome translocations specific for the myeloma cell clone: a new type of cancer–host partnership? The Journal of Pathology, 2007. 211(1): p. 10–17.

11. Tan, C.-C., et al., Breast cancer cells obtain an osteomimetic feature via epithelial-mesenchymal transition that have undergone BMP2/RUNX2 signaling pathway induction. Oncotarget, 2016. 7.

12. Awolaran, O., S.A. Brooks, and V. Lavender, Breast cancer osteomimicry and its role in bone specific metastasis; an integrative, systematic review of preclinical evidence. The Breast, 2016. 30: p. 156–171.

13. Gu, C., et al., Targeting initial tumour–osteoclast spatiotemporal interaction to prevent bone metastasis. Nature Nanotechnology, 2024. 19(7): p. 1044–1054.

14. Wan, Y., et al., Mechanical control of osteoclast fusion by membrane-cortex attachment and BAR proteins. Journal of Cell Biology, 2025. 224(7): p. e202411024.

15. Ma, Q., et al., Osteoclast-derived apoptotic bodies couple bone resorption and formation in bone remodeling. Bone Research, 2021. 9(1): p. 5.

16. Harre, U., et al., Moonlighting osteoclasts as undertakers of apoptotic cells. Autoimmunity, 2012. 45(8): p. 612–619.

17. Pekowska, A., et al., A unique H3K4me2 profile marks tissue-specific gene regulation. Genome Res, 2010. 20(11): p. 1493–502.

18. Arellano, D.L., et al., Bone Microenvironment-Suppressed T Cells Increase Osteoclast Formation and Osteolytic Bone Metastases in Mice. Journal of Bone and Mineral Research, 2022. 37(8): p. 1446–1463.

19. Shiirevnyamba, A., et al., Enhancement of osteoclastogenic activity in osteolytic prostate cancer cells by physical contact with osteoblasts. British Journal of Cancer, 2011. 104(3): p. 505–513.

20. Ding, K., et al., Single cell heterogeneity and evolution of breast cancer bone metastasis and organoids reveals therapeutic targets for precision medicine. Annals of Oncology, 2022. 33.

21. Le Pape, F., G. Vargas, and P. Clézardin, The role of osteoclasts in breast cancer bone metastasis. Journal of Bone Oncology, 2016. 5(3): p. 93–95.

22. Aguirre, L.A., et al., Tumor stem cells fuse with monocytes to form highly invasive tumor-hybrid cells. Oncoimmunology, 2020. 9(1): p. 1773204.

23. Chou, C.-W., et al., Phagocytosis-initiated tumor hybrid cells acquire a c-Myc-mediated quasi-polarization state for immunoevasion and distant dissemination. Nature Communications, 2023. 14.

24. Madel, M.-B., et al., Immune Function and Diversity of Osteoclasts in Normal and Pathological Conditions. Frontiers in Immunology, 2019. 10(1408).

25. Raymaekers, K., et al., The vasculature: a vessel for bone metastasis. Bonekey Rep, 2015. 4: p. 742.

26. Kolb, A.D. and K.M. Bussard, The Bone Extracellular Matrix as an Ideal Milieu for Cancer Cell Metastases. Cancers (Basel), 2019. 11(7).

27. Jiang, Y., et al., Targeting extracellular matrix stifness and mechanotransducers to improve cancer therapy. J Hematol Oncol, 2022. 15(1): p. 34.

28. Sugatani, T. and K. Hruska, Akt1/Akt2 and Mammalian Target of Rapamycin/Bim Play Critical Roles in Osteoclast Diferentiation and Survival, Respectively, Whereas Akt Is Dispensable for Cell Survival in Isolated Osteoclast Precursors. The Journal of biological chemistry, 2005. 280: p. 3583–9.

29. Lv, L. and X. Zhou, Targeting Hippo signaling in cancer: novel perspectives and therapeutic potential. MedComm (2020), 2023. 4(5): p. e375.

30. Furesi, G., M. Rauner, and L.C. Hofbauer, Emerging Players in Prostate Cancer&#x2013;Bone Niche Communication. Trends in Cancer, 2021. 7(2): p. 112–121.

31. Wang, H., et al., NeuroD4 converts glioblastoma cells into neuron-like cells through the SLC7A11-GSH-GPX4 antioxidant axis. Cell Death Discovery, 2023. 9(1): p. 297.

32. Wechman, S.L., et al., Vascular mimicry: Triggers, molecular interactions and in vivo models. Adv Cancer Res, 2020. 148: p. 27–67.

33. Lovero, D., et al., Correlation between targeted RNAseq signature of breast cancer CTCs and onset of bone-only metastases. British Journal of Cancer, 2022. 126(3): p. 419–429.

34. Takeshita, S., K. Kaji, and A. Kudo, Identification and Characterization of the New Osteoclast Progenitor with Macrophage Phenotypes Being Able to Diferentiate into Mature Osteoclasts*. Journal of Bone and Mineral Research, 2009. 15(8): p. 1477–1488.

35. Clohisy, D.R., et al., Human breast cancer induces osteoclast activation and increases the number of osteoclasts at sites of tumor osteolysis. J Orthop Res, 1996. 14(3): p. 396–402.

36. Kuchimaru, T., et al., A reliable murine model of bone metastasis by injecting cancer cells through caudal arteries. Nature Communications, 2018. 9(1): p. 2981.

37. Andrews, S., FastQC: A quality control tool for high throughput sequence data.

38. Dobin, A., et al., STAR: ultrafast universal RNA-seq aligner. Bioinformatics, 2013. 29(1): p. 15–21.

39. Li, B. and C.N. Dewey, RSEM: accurate transcript quantification from RNA-Seq data with or without a reference genome. BMC Bioinformatics, 2011. 12: p. 323.

40. Love, M.I., W. Huber, and S. Anders, Moderated estimation of fold change and dispersion for RNA-seq data with DESeq2. Genome Biol, 2014. 15(12): p. 550.

41. Wu, T., et al., clusterProfiler 4.0: A universal enrichment tool for interpreting omics data. Innovation (Camb), 2021. 2(3): p. 100141.

42. Wickham, H., ggplot2: Elegant Graphics for Data Analysis. Springer-Verlag New York, 2016. 978-3-319-24277-4.

43. Hao, Y., et al., Dictionary learning for integrative, multimodal and scalable single-cell analysis. Nature Biotechnology, 2024. 42(2): p. 293–304.

44. Hänzelmann, S., R. Castelo, and J. Guinney, GSVA: gene set variation analysis for microarray and RNA-Seq data. BMC Bioinformatics, 2013. 14(1): p. 7.

