## Supplementary material for "Breast cancer interactions with osteoclasts generate osteoclast-tumor hybrid-like cells through dynamic non-canonical cell fusion and cell-in-cell processes": Supp. Table 1

**Supplementary Table 1. Primer sequences used for quantitative RT–PCR**

| Gene symbol | Forward primer (5′–3′) | Reverse primer (5′–3′) |
|---|---|---|
| Spi1 | TTACAGGCGTGCAAAATGGAA | GACGTTGGTATAGCTCTGAATCG |
| Fcer1g | ATCTCAGCCGTGATCTTGTTCT | ACCATACAAAAACAGGACAGCAT |
| Dap12 (Tyrobp) | CCCAAGATGCGACTGTTCTTC | GTCCCTTGACCTCGGGAGA |
| Dcstamp | GGGGACTTATGTGTTTCCACG | ACAAAGCAACAGACTCCCAAAT |
| Ocstamp | CTGTAACGAACTACTGACCCAGC | CCAGGCTTAGGAAGACGAAGA |
| Trem2 | CTGGAACCGTCACCATCACTC | CGAAACTCGATGACTCCTCGG |
| Car2 | GATAAAGCTGCGTCCAAGAGC | GCATTGTCCTGAGAGTCATCAAA |
| Atp6v0d2 | CTGGTTCGAGGATGCAAAGC | GTTGCCATAGTCCGTGGTCTG |
| Ripk3 | GTGCTACCTACACAGCTTGAAC | CCCTCCCTGAAACGTGGAC |
| Oscar | CCTAGCCTCATACCCCCAG | CGTTGATCCCAGGAGTCACAA |
| Ctsk | CTCGGCGTTTAATTTGGGAGA | TCGAGAGGGAGGTATTCTGAGT |
| Cd74 | AGATGCGGATGGCTACTCC | TCATGTTGCCGTACTTGGTAAC |
| Fos | CGGGTTTCAACGCCGACTA | TGGCACTAGAGACGGACAGAT |
| Mmp9 | GCAGAGGCATACTTGTACCG | TGATGTTATGATGGTCCCACTTG |
| Syna | ATGGTTCGTCCTTGGGTTTTC | GTGTTGAGTGAGGTTTACCAGG |
| Acp5 | CACTCCCACCCTGAGATTTGT | CCCCAGAGACATGATGAAGTCA |
| Mmp2 | ACCTGAACACTTTCTATGGCTG | CTTCCGCATGGTCTCGATG |
